# Cortical Presynaptic Boutons Progressively Engulf Spinules As They Mature

**DOI:** 10.1101/2020.07.14.202663

**Authors:** Charles Campbell, Sarah Lindhartsen, Adam Knyaz, Alev Erisir, Marc Nahmani

## Abstract

Despite decades of discussion in the neuroanatomical literature, the role of the synaptic ‘spinule’ in synaptic development and function remains elusive. Canonically, spinules are finger-like projections that emerge from postsynaptic spines and can become enveloped by presynaptic boutons. When a presynaptic bouton encapsulates a spinule in this manner, the membrane apposition between the spinule and surrounding bouton can be significantly larger than the membrane interface at the synaptic active zone. Hence, spinules may represent a mechanism for extrasynaptic neuronal communication, and/or may function as structural ‘anchors’ that increase the stability of cortical synapses. Yet despite their potential to impact synaptic function, we have little information on the percentages of developing and adult cortical bouton populations that contain spinules, the percentages of these cortical spinule-bearing boutons (SBBs) that contain spinules from distinct neuronal/glial origins, or whether the onset of activity or cortical plasticity are correlated with increased prevalence of cortical SBBs. Here, we employed 2D and 3D electron microscopy to determine the prevalence of spinules in excitatory presynaptic boutons at key developmental time points in the primary visual cortex (V1) of female and male ferrets. We find that the prevalence of SBBs in V1 increases across postnatal development, such that ~25% of excitatory boutons in late adolescent ferret V1 contain spinules. In addition, we find that a majority of spinules within SBBs at later developmental time points emerge from postsynaptic spines and adjacent boutons/axons, suggesting that synaptic spinules may enhance synaptic stability and allow for axo-axonal communication in mature sensory cortex.

**Significance Statement:** Synaptic spinules are finger-like projections from neurites that can become completely embedded within presynaptic boutons, potentially enhancing synaptic communication and stability. Yet while their existence has been discussed for decades, spinule prevalence, projection origins, and relationship to neocortical sensory activity remain unknown. In this study, we employed 2D and 3D electron microscopy techniques to characterize the development of excitatory cortical spinule-bearing boutons (SBBs) and their relationship to sensory activity and plasticity. Our results demonstrate that neocortical presynaptic SBB prevalence is not correlated with the onset of sensory activity or heightened cortical plasticity, and that by late adolescence nearly one-quarter of presynaptic boutons contain a spinule. Hence, synaptic spinules may play progressively important roles in cortical function as these synapses mature.

## Introduction

Synapses underlie the sensory, motor, and cognitive processing functions of nervous systems and alterations of synaptic communication and stability directly affect animal behavior and memory (Sherrington, 1906; Liewald et al., 2008; Liu et al., 2012). Moreover, changes in synaptic morphology can accurately predict changes in synaptic strength and stability (Murthy et al., 2001; Ostroff et al., 2002; Branco et al., 2010; Holderith et al., 2012; Araya et al., 2014; Meyer et al., 2014; Quinn et al., 2019). Yet, a ubiquitous and potentially crucial synaptic structure, the synaptic spinule, remains enigmatic and underexplored.

Spinules are components of synapses that are conserved across the animal kingdom (Case et al., 1972; Bailey et al., 1979) yet have an unknown function. Canonically, spinules are thin, finger-like invaginating projections from postsynaptic dendritic spines that can become enveloped by other neurites (Pappas and Purpura, 1961; Westrum and Blackstad, 1962; Tarrant and Routtenberg, 1977; Spacek and Harris, 2004). When a neuronal structure such as a presynaptic bouton envelops a spinule, the membrane interface between the spinule-projecting structure and the enveloping bouton increases, in some cases up to 26 times the size of a synapse’s ‘active zone’ (Rodriguez-Moreno et al., 2018). Thus, spinules may represent an undescribed mechanism for neuronal communication, and/or they may function as structural ‘anchors’ that increase synaptic strength and stability. If true, an increase in spinule-bearing synapses within a functionally defined microcircuit could enable large changes in brain function, such as increased memory retention or improved recovery after injury.

To date, the best anatomically and physiologically characterized form of spinules are those of the mammalian hippocampus, specifically spinules emanating from the postsynaptic spines of CA1 stratum radiatum and dentate gyrus pyramidal cell dendrites in adult rat (Tarrant and Routtenberg, 1977; Geinisman et al., 1994; Toni et al., 1999; Spacek and Harris, 2004). While spinules often originate from spines and become engulfed by presynaptic boutons, the term ‘spinules’ has broadened to describe any projection from a neurite or glia that becomes enveloped within another neuronal or glial process (Petralia et al., 2015). In CA1 stratum radiatum, ~32% of spines project spinules, and most of these spinules are enveloped by presynaptic boutons (~90%) (Spacek and Harris, 2004). Intriguingly, the percentage of CA1 spines displaying spinules increases following increases in neuronal activity, such as electrical and chemical long-term potentiation (LTP) protocols (Geinisman et al., 1994; Toni et al., 1999; Harris et al., 2003; Ueda and Hayashi, 2013), high extracellular K_+_ (Tao-Cheng et al., 2009), and estradiol receptor activation (Murphy and Andrews, 2000). These data have led to the suggestion that spinules function as activity-dependent neuronal circuit remodeling and signaling elements throughout the brain (Spacek and Harris, 2004), or as a mechanism of presynaptic membrane retrieval during periods of heightened activity (Tao-Cheng et al., 2009).

Yet, whether these results from hippocampus generalize to other areas of the brain (e.g. neocortex) remains unclear, and whether heightened levels of developmental activity/plasticity correlate with heightened levels of spinule prevalence in vivo remains unknown. Furthermore, few studies have investigated the percentages of presynaptic boutons that contain spinules, a key component in understanding the potential for spinule-induced synaptic communication and/or stability. For example, a 2D ultrastructural study of thalamocortical (TC) boutons found that 28% of TC boutons in layer 4 (L4) of primary visual cortex (V1) contained spinule-like (i.e. putative spinules) 2D cross-sections (Erisir and Dreusicke, 2005), and a 3D study of TC boutons in L4 of barrel cortex reported that 13% of reconstructed postsynaptic dendritic spines projected a spinule into TC boutons (Rodriguez-Moreno et al., 2018). Thus, spinules may be a prominent feature of neocortical synapses.

Here, we used the broader definition of ‘spinules’ to describe invaginating projections from neurites or glia that are enveloped by excitatory cortical presynaptic boutons. We sought to determine the proportion of synaptic excitatory spinule-bearing boutons (SBBs) within the excitatory bouton population in V1 across postnatal development, and the origins of spinules projecting into these SBBs. In addition, we examined whether the onset of sensory activity or heightened levels of cortical plasticity correlated with a commensurate increase in the proportion of SBBs in V1. Using 2D and 3D electron microscopy analyses, we find that: (1) the proportion of SBBs increases in parallel with the developmental maturation of excitatory boutons and strengthening of synapses in V1, (2) that SBBs preferentially engulf spinules from postsynaptic spines and adjacent boutons/axons as they mature, and that (3) ~25% of the excitatory synaptic boutons in late adolescent ferret contain spinules.

## Materials and Methods

### Animals

A total of ten male and female ferrets were used for this study, postnatal day (p)21 (n=1), p28 (n=1), p46 (n=1), p47 (n=1), p60 (n=2), p66 (n=1), >p90 (n=3). These ages were chosen because they are key developmental time points: before eye opening (i.e. before ~p32), the peak of the critical period for ocular dominance and thalamocortical axon morphological plasticity (~p46), the end of this critical period in ferret binocular V1 (~p60), and ages approaching ferret sexual maturity (>p90) (Issa et al., 1999). For the purposes of 2D analyses, measurements from animals at similar developmental time points were grouped (i.e. p21-28 (n=2), p46-47 (n=2), p60-66 (n=3), and >p90 (n=3)), after determining that intragroup (e.g. p21 vs. p28) measurements of synapse length and bouton cross-section area were not statistically significant (One-Way ANOVA and post-hoc Bonferroni corrected t Tests, p>0.1; data not shown). All animal procedures and protocols were in accordance with NIH guidelines for humane handling of animals and were approved by the Institutional Animal Care and Use Committee at the University of Virginia.

### Perfusions

Animals were given an overdose of Nembutal (in excess of 50 mg/kg) and were perfused transcardially with 4 % paraformaldehyde and 0.5% glutaraldehyde in 0.1 M phosphate buffer (PB; pH 7.4). To prevent the reported increase in spinule prevalence due to longer perfusion times (Tao-Cheng et al., 2009), we only included animals in this study where perfusion time from thorax incision to fixation was < 100 seconds. Following perfusions, brains were removed and kept in 4% paraformaldehyde overnight at 4 °C. On the following day, each brain was dissected, and occipital lobe blocks were placed in a vibratome and coronal sections of V1 were cut at 60 μm. Floating coronal sections were immediately treated with 1 % sodium borohydride, rinsed 5-6 times in phosphate-buffered saline (PBS), and free-floating sections were stored in PBS containing 0.05% sodium azide at 4°C.

### Transmission Electron Microscopy

Sections prepared for transmission electron microscopy (TEM) examination were rinsed in 0.1 M PB and then immersed in 1% osmium tetroxide (in 0.1 M PB) for 1 hour. Next, sections were rinsed in 0.1 M PB, dehydrated in a series of ethanol dilutions, and then incubated in 4% uranyl acetate (in 70% ethanol) at 4 °C overnight. Sections were then dehydrated in acetone and incubated in a series of three progressively concentrated acetone-Epon 812 resin (EMS, Cat#RT14120) mixtures, each for 4 hours – overnight at room temperature. Next, sections were flat embedded between two pieces of plastic film (EMS, Cat#50425-10) and placed in a 60 °C oven overnight. Areas within layer 4 of the binocular region of V1 of embedded coronal sections were marked under light microscopy using established white matter landmarks (Law et al., 1988), excised from the flat embed, placed in a capsule (EMS, Cat#70010-B), and filled with Epon 812 resin. Layer 4 of binocular V1 is a site of robust physiological and morphological developmental plasticity (Issa et al., 1999; Coleman et al., 2010; Maffei et al., 2010). Next, tissue blocks were cured in a 60 °C oven for 36-48 hours or until polymerized. Outlines and landmarks of layer 4 of binocular V1 were then drawn using a camera lucida and this area of interest was cut into a trapezoid on the EPON block. Ultrathin 60-70 nm thick sections were then cut using an ultramicrotome (Leica Ultracut, Leica Microsystems) and collected on TEM copper and nickel mesh grids. Ultrathin sections were viewed using a Philips CM 100 or JEOL JEM 1400 TEMs equipped with a 2K x 3k Olympus Morada and a 2K x 2K Gatan Ultrascan 1000XP camera, respectively. Digital electron micrographs were taken at final magnifications on these TEMs to achieve ~0.5 nm / pixel resolution. Potential tissue shrinkage due to TEM/FIBSEM tissue processing was not expected to differentially affect asymmetric synapses across age groups (Korogod et al., 2015) and was not corrected for in our analyses.

Ultrathin sections were selected at random from each TEM grid and nonoverlapping TEM images were taken across L4 within each ultrathin section. To ensure that we did not image the same synapse more than once, imaged ultrathin sections at each age were taken ≥ 2 μm apart in depth (z), which was ~ 3.5 times greater than the depth of the largest postsynaptic density from our FIBSEM analyses. In each ultrathin section we analyzed every Gray’s Type I ‘asymmetric’ excitatory synapse (Gray, 1959) we encountered, being careful to locate and include smaller synapses within these sections in our analyses. Putative excitatory synapses were only included in our analyses if they (1) had parallel alignment of presynaptic and postsynaptic membranes at the active zone, (2) had ≥ 3 presynaptic vesicles, and (3) had a prominent asymmetric postsynaptic density opposite these presynaptic vesicles. All excitatory synapses meeting these criteria were analyzed using FIJI (Schindelin et al., 2012) software to record their bouton cross section areas, synapse cross section lengths, and spinule cross section areas (if present).

In 2D TEM images, spinules were identified as double membrane-bound structures (i.e. inner spinule lipid bilayer and outer bouton lipid bilayer) within excitatory synaptic boutons. In addition, spinules were only included in our analyses if they were ≥ 2.5 times the size (diameter of shortest spinule axis) of the averaged sized synaptic vesicle at that age, and if their interiors were electron translucent. We adopted these conservative criteria in order to differentiate spinules from synaptic vesicles, elongated endoplasmic reticula (Wu et al., 2017), and relatively electron opaque lysosomes (Marty and Peschanski, 1994) in our 2D TEM analyses. In the event that multiple spinule cross sections were present within a single SBB, these spinule cross section areas were summed for the purposes of our spinule cross section area analyses.

### Focused Ion Beam Scanning Electron Microscopy

Tissue blocks from a p21, p46, p60, and >p90 animal, originally used in our 2D TEM analyses, were retrimmed and prepared for focused ion beam scanning electron microscopy (FIBSEM). First, a series of 60 nm ultrathin sections were cut to a depth of ~ 3 μm from each block to ensure we would be analyzing novel synapses in our 3D analyses. All processing and imaging of FIBSEM blocks was performed at the Multiscale Microscopy Core at Oregon Health and Sciences University. Tissue blocks were mounted onto standard 12.5-mm flat SEM stubs (Ted Pella, Cat#16111) using Leitsilber 200 silver paint (Ted Pella, Cat#16035). The resulting blocks were then trimmed using a Trim90 diamond knife (Diatome) and the final sample blocks were coated with 8 nm of carbon using a Leica ACE 600 unit. Regions of interest from the block faces were imaged utilizing Thermo Fisher’s Auto Slice & View™ software package to automate the serial sectioning and data collection processes. Each block face was scanned on a FEI Helios 660 DualBeam SEM at a 52° tilt and with a 4.2 mm working distance. For each slice, 4 nm of resin was removed at 30 keV and 0.79 nA. Each slice was imaged with a 3 kV acceleration voltage and 400 pA current in backscatter mode with inverse contrast using the in-column detector. Images of 6144 × 4096 pixels were acquired at a resolution of 4 nm/pixel.

We obtained aligned 4 nm isotropic FIBSEM image stacks from a p21, p46, p60, and >p90 animal. Image stacks of L4 V1 were of the following volumes: p21 (15.1 × 14.1 × 2.8 μm), p46 (9.7 × 8.4 × 2.7 μm), p60 (24.2 × 16.2 × 2.4 μm), and >p90 (24.2 × 16.2 × 2.4 μm). All serial image stacks were analyzed in FIJI. We located every excitatory synapse meeting our excitatory synapse criteria within each image stack and analyzed them in 3D for (1) the presence/absence of a spinule, (2) presence/absence of a perforated postsynaptic density, and (3) the origin of the spinule (if present). Importantly, spinules were only included in our FIBSEM analyses if they could be observed invaginating into a presynaptic bouton and were then observed as membrane-bound structures (i.e. unambiguous visualization of lipid bilayer of spinule surrounded by bouton’s lipid bilayer) within the presynaptic bouton in at least one image. This additional positive spinule identification criteria necessarily differs from our 2D analysis criteria, since 4 nm FIBSEM z-resolution allows one to visualize a spinule’s invagination site into each SBB. In contrast, our 2D analyses required stricter size criteria to identify spinules from 2D profiles (i.e. ≥ 2.5 times a synaptic vesicle), due to a lack of information on the potential origins of membrane-bound objects within 2D TEM images. Therefore, our 2D analyses may have excluded smaller spinules and consequently yielded reduced spinule prevalence. Spinule origins in FIBSEM stacks were determined by tracking them within the image volume to their parent neurite/glia. When it was not possible to identify the parent neurite/glia from which a spinule emerged (e.g. due to section loss/defect, or an object lacking one or more identifiable morphological criteria by the end of the image volume) we assigned this spinule to the “unknown” origin category. Following quantification of spinule origins, we reconstructed SBBs from representative parent origins, including their synaptic partners and spinule parent neurites/glia, using Reconstruct software (Fiala, 2005). 3D reconstruction surfaces were then remeshed and rendered semi-transparent for spinule and PSD visualization using Blender open-source software.

### Statistics

Descriptive statistics for all group comparisons in this study are listed in Table 3. For intragroup comparisons of spinule area, bouton area, and synapse length cross sections within developmental age groups, we performed One-Way ANOVAs (p60-p66 and >p90 age groups) and post-hoc Bonferroni corrected t Tests (p21-28 and p46-47 age groups) to compare means from animals within each group and found that there were no significant differences (p>0.1) between animals within any single age group (data not shown). Accordingly, we grouped 2D data within each age group into a single group mean. To compare spinule area, bouton area, and synapse length cross sections between developmental groups, we then performed one-way ANOVAs, followed by Bonferroni post hoc two-tailed t Tests if our ANOVA results were significant at the p < 0.05 level (Bonferroni correction set at p < 0.0168). To compare spinule prevalence within 2D TEM images between age groups, we used two-tailed Chi-Square tests with Bonferroni correction (4 x 2 contingency table, three comparisons, Bonferroni corrected significance level set at p < 0.0168). For our Gardner-Altman estimation statistic plots (Gardner and Altman, 1986), morphological measurement distributions were compared using non-parametric Mann-Whitney U tests.

For FIBSEM analyses, to compare the proportions of SBBs, perforated PSDs, and spinule origins between animals, we used two-tailed Chi-Square tests with Bonferroni correction (4 x 2 contingency table, three comparisons, Bonferroni corrected significance level set at p < 0.0168). For all figures, *** = p < 0.001, ** = p < 0.015, and * = p < 0.05.

## Results

To determine whether the onset of sensory activity or cortical plasticity state correlate with spinule-bearing bouton (SBB) prevalence (i.e. the number of SBBs divided by the total number of sampled excitatory presynaptic boutons), we analyzed excitatory synaptic boutons within L4 from ferret V1 across key developmental stages. We reasoned that if the onset of sensory activity and/or heightened cortical plasticity is primarily responsible for increasing the proportion of SBBs in V1, SBB prevalence should increase rapidly between when V1 cells first receive sensory activity (i.e. eye-opening) and when Hebbian and homeostatic plasticity are maximally expressed in V1 (approximately postnatal day (p)32 – p46 in ferret) (Issa et al., 1999). In contrast, if the presence of a spinule within a presynaptic bouton is related to its physiological maturation and morphological stability (Cheetham and Fox, 2010; Smith et al., 2015; Qiao et al., 2016), we expect to see an increase in SBB prevalence in parallel with this progressive functional maturation. To this end, we prepared sections from L4 of binocular V1 for transmission electron microscopy (TEM) from ferrets at key postnatal ages: p21-28 (before eye opening; n = 2 animals), p46-47 (height of plasticity in V1; n = 2 animals), p60-66 (end of critical period in V1; n = 3 animals), and >p90 (ferrets between p90 – p150 that were nearing sexual maturity; n= 3 animals). We took non-overlapping 2D TEM images within L4 from animals within each of these age groups and quantified morphological features correlated with synaptic strength (Murthy et al., 2001; Ostroff et al., 2002; Branco et al., 2010; Holderith et al., 2012; Araya et al., 2014; Meyer et al., 2014), as well as the presence or absence of SBBs, at every excitatory synapse we encountered (see Methods).

### Morphometry of Excitatory Synaptic Boutons and Spinules Across Development

Qualitatively, p21-p28 excitatory synapses had less pronounced postsynaptic densities (PSDs) and often displayed *en passant* bouton morphology (**Fig. 1**), whereas at older ages PSDs appeared to become larger and thicker and excitatory boutons displayed more varied and complex morphologies. Indeed, while our quantitative analyses revealed that 2D synapse length cross sections did not appreciable change from p21 to p66, PSD lengths did show a significant 16% increase between p60-66 and >p90 ages (**Fig. 2A, Tables 1 & 3**). Cortical synaptic boutons have been shown to increase in size over postnatal development (Erisir and Dreusicke, 2005). Yet interestingly, we found that bouton cross section areas were largest at p21-28 (before eye opening), dropped by 93% at p46-47, and then increased significantly until >p90 (**Fig. 2B**). This initial developmental decrease in bouton areas has been reported previously for excitatory synaptic boutons in L4 (Dufour et al., 2016), and may stem from a preponderance of elongated *en passant* boutons at early developmental ages. Following this initial decrease however, excitatory boutons in L4 showed a steady, nearly linear increase in their cross-section areas from p46 to >p90 (**Fig. 2B, Fig. 2-1, Tables 1 & 3**).

**Table 1.** Means, Standard Error, and Sample Sizes for 2D TEM Morphology Analyses. Means and standard error of the mean for individual animals and developmental age group are listed for each 2D TEM analysis measurement (column titles). Population sizes are listed for individual animals and age groups under “Total Boutons” and “Total Synapses” column. PSD = postsynaptic density; SBBs = spinule-bearing boutons; Spinule:Bouton Area % = percentage of bouton area occupied by spinule area.

|  | Age | Total Boutons | Total Synapses | PSD Length (μm) | Bouton Area (um<sup>2</sup>) | % SBBs | Spinule Area (um<sup>2</sup>) | Spinule:Bouton Area (%) |
| --- | --- | --- | --- | --- | --- | --- | --- | --- |
|  | p21 | 70 | 74 | 0.29 | 0.34 | 4% | 0.04 | 12% |
|  | p28 | 71 | 78 | 0.27 | 0.42 | 7% | 0.02 | 6% |
| **Group Sum** |  | **141** | **152** |  |  |  |  |  |
| **Group Mean ± S.E.M. (n = animals)** |  |  |  | 0.28 ± 0.01 | 0.38 ± 0.03 | 6 ± 1.4% | 0.03 ± 0.01 | 9 ± 3.1% |
| **Cumulative Mean ± S.E.M. (n = sampled structures)** |  |  |  | **0.28 ± 0.01** | **0.38 ± 0.03** | **6 ± 1.4%** | **0.03 ± 0.01** | **6 ± 1.5%** |
|  | p46 | 102 | 102 | 0.28 | 0.19 | 6% | 0.02 | 6% |
|  | p47 | 94 | 94 | 0.29 | 0.21 | 6% | 0.03 | 12% |
| **Group Sum** |  | **196** | **196** |  |  |  |  |  |
| **Group Mean ± S.E.M. (n = animals)** |  |  |  | 0.29 ± 0.01 | 0.20 ± 0.01 | 6 ± 0.3% | 0.03 ± 0.001 | 9 ± 3.2% |
| **Cumulative Mean ± S.E.M. (n = sampled structures)** |  |  |  | **0.29 ± 0.01** | **0.20 ± 0.01** | **6 ± 0.3%** | **0.02 ± 0.004** | **8 ± 3.2%** |
|  | p60_1 | 25 | 29 | 0.28 | 0.21 | 10% | 0.04 | 14% |
|  | p60_2 | 111 | 113 | 0.29 | 0.28 | 13% | 0.01 | 4% |
|  | p66_1 | 118 | 124 | 0.31 | 0.28 | 11% | 0.04 | 14% |
| **Group Sum** |  | **254** | **266** |  |  |  |  |  |
| **Group Mean ± S.E.M. (n = animals)** |  |  |  | 0.30 ± 0.01 | 0.26 ± 0.02 | 11 ± 0.8% | 0.03 ± 0.01 | 11 ± 3.4% |
| **Cumulative Mean ± S.E.M. (n = sampled structures)** |  |  |  | **0.30 ± 0.01** | **0.27 ± 0.01** | **11 ± 0.8%** | **0.03 ± 0.01** | **10 ± 1.7** |
|  | >p90_1 | 61 | 61 | 0.32 | 0.29 | 16% | 0.06 | 16% |
|  | >p90_2 | 149 | 150 | 0.36 | 0.33 | 15% | 0.02 | 5% |
|  | >p90_3 | 109 | 112 | 0.34 | 0.35 | 22% | 0.04 | 4% |
| **Group Sum** |  | **319** | **323** |  |  |  |  |  |
| **Group Mean ± S.E.M. (n = animals)** |  |  |  | 0.35 ± 0.01 | 0.32 ± 0.01 | 18 ± 2.1% | 0.04 ± 0.01 | 8 ± 4% |
| **Cumulative Mean ± S.E.M. (n = sampled structures)** |  |  |  | **0.35 ± 0.01** | **0.33 ± 0.01** | **18 ± 2.1 %** | **0.04 ± 0.01** | **8 ± 1%** |

**Figure 1.**
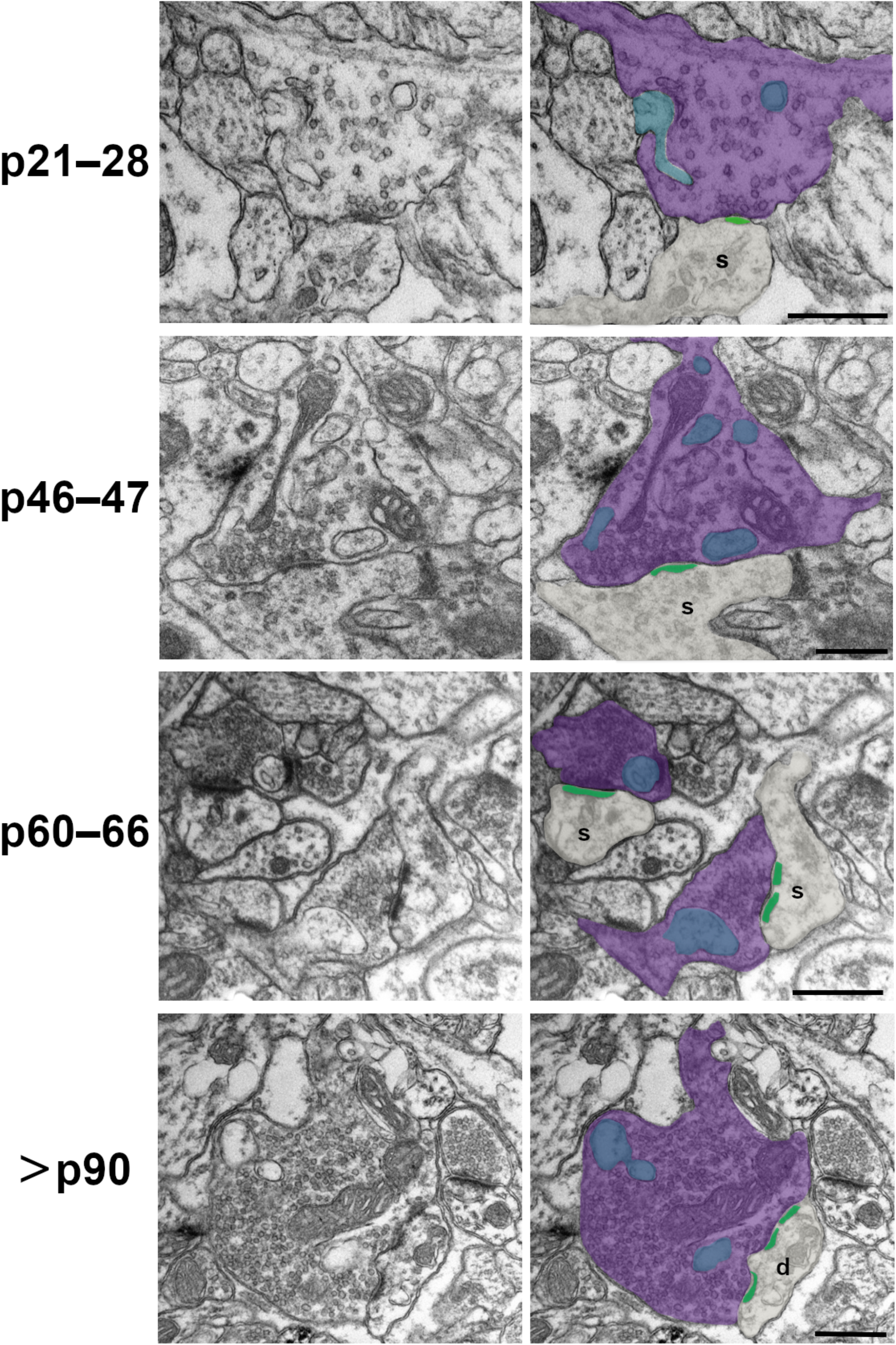
Development of Excitatory Spinule-Bearing Bouton (SBB) Morphology. Spinules (blue) were observed within cortical boutons (purple) at every postnatal (p) age examined. In 2D TEM sections, spinules were occasionally observed invaginating into SBB cross sections (e.g. p21-28), however most spinules appeared as ovoid or circular double membrane-bound structures encapsulated within their ‘host’ bouton. Postsynaptic densities (green) at SBB synapses sometimes contained perforations (e.g. p60-66 and >p90 panels). Note that SBBs at times displayed multiple spinule cross sections, yet in our 2D TEM analyses it was not possible to attribute these to single spinules with complex morphologies, or to multiple spinules protruding into a single SBB. s = postsynaptic spine; d = postsynaptic dendrite; Scale bars = 0.5 μm.

**Figure 2.**
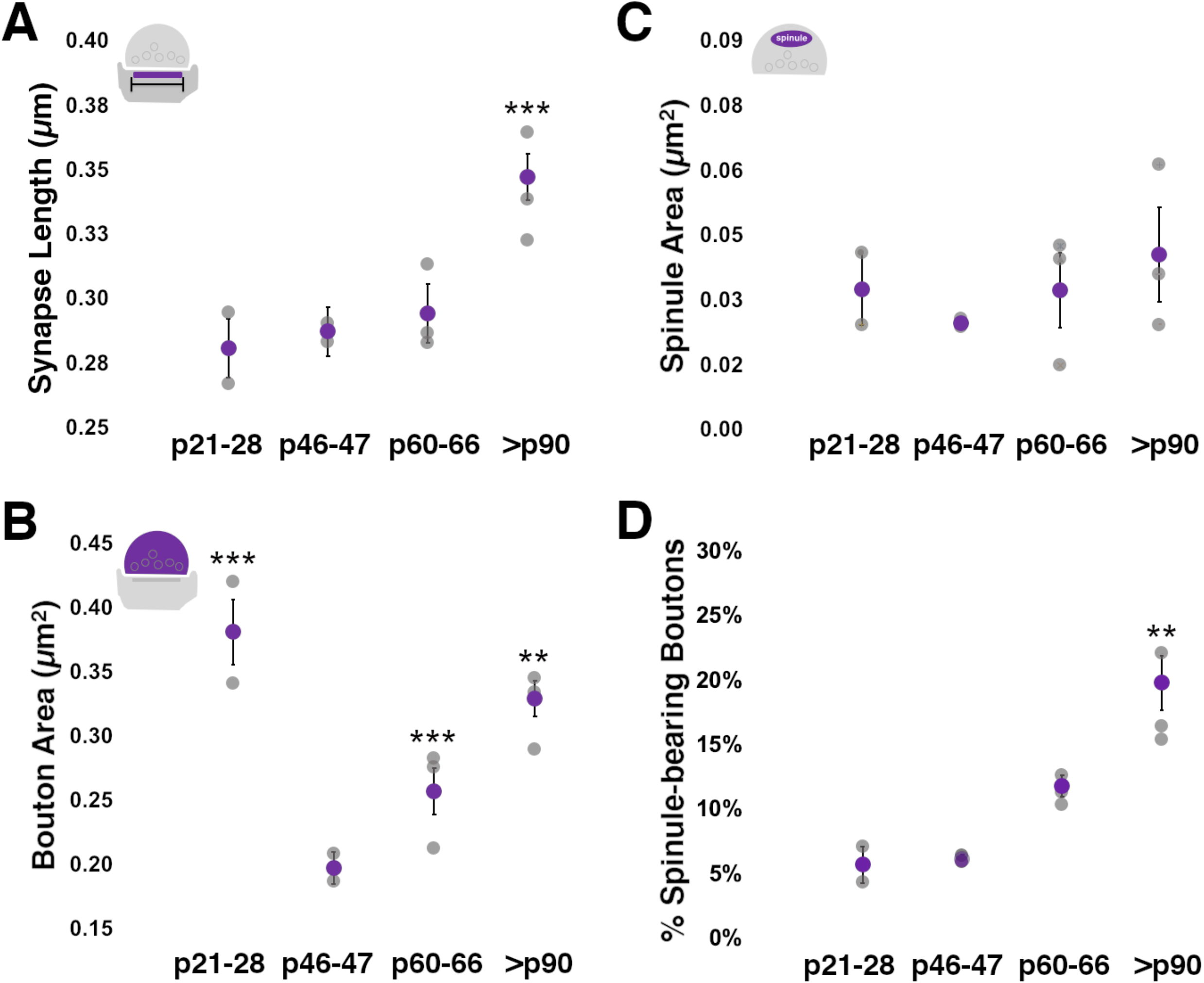
Excitatory Bouton Areas and Spinule Prevalence Increase Over Development. **A**. 2D synapse cross section length, as defined by the postsynaptic density (PSD), remains relatively stable before eye-opening (p28) and the end of the critical period for plasticity in ferret visual cortex (p60). However, PSD length increases substantially from the end of the critical period until at least the cusp of maturity (>p90 ages). **B**. Excitatory presynaptic bouton cross section areas are largest at a time when most boutons appear to have enlarged en-passant morphology (p21-28). Bouton size decreases significantly by p46-47 and then shows a steady increase until at least p90. **C**. Average spinule cross section areas do not appreciable change across the postnatal ages examined. Note that spinule areas show relatively large inter-animal variation attributed to the variation in spinule sizes from distinct parent neurite/glia origins. **D**. Excitatory spinule-bearing boutons (SBBs) are most prevalent in late adolescence versus at the end of critical period (p60-66) in ferret primary visual cortex. A trend toward increased SBB prevalence is seen between p46-47 and >90 ages. Gray circles = individual animal means; Purple circles = group means ± S.E.M. ** = p < 0.015; *** = p < 0.001.

**Figure 2-1.**
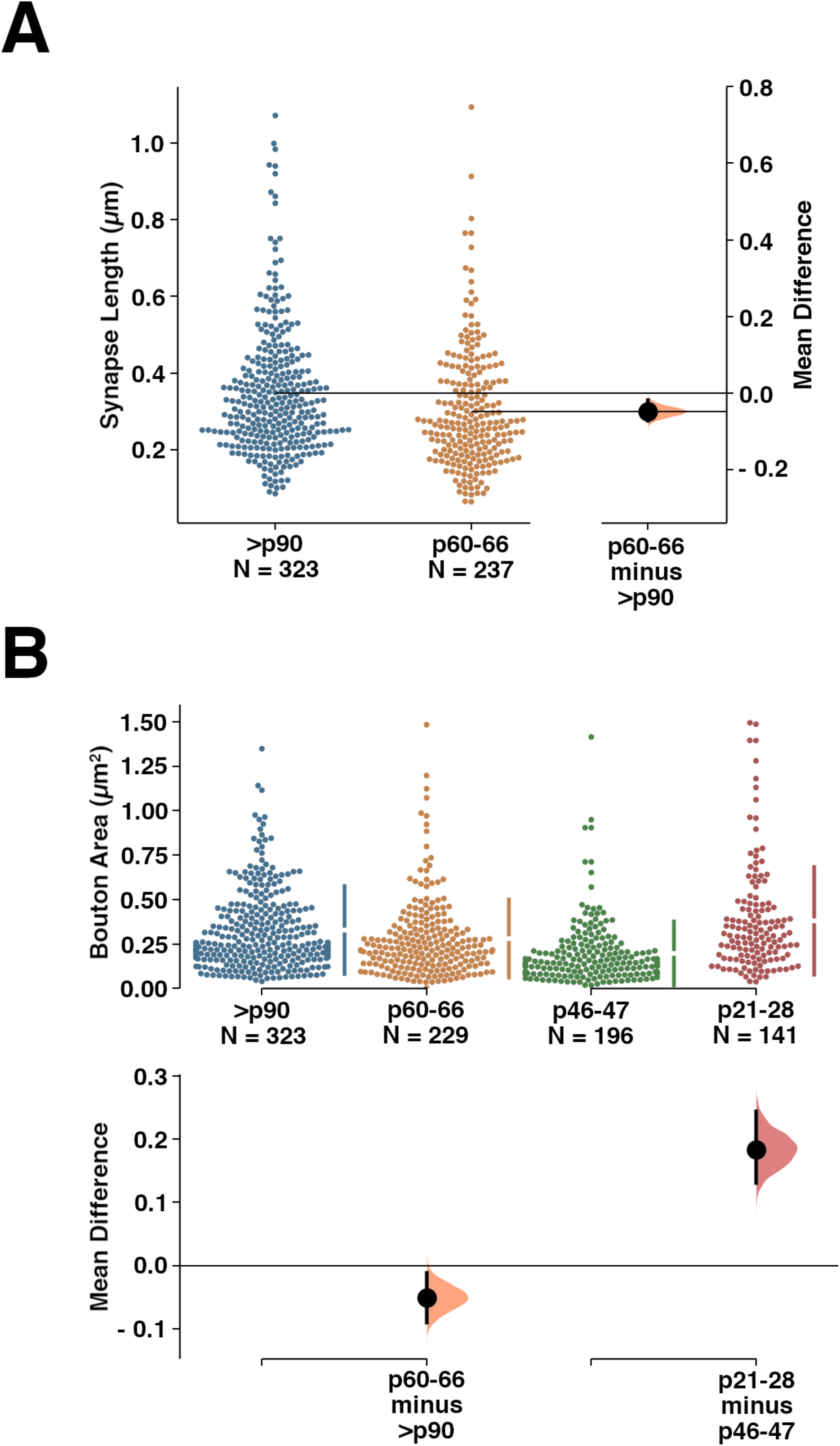
Estimation Statistics Plots Showing Mean Differences Between Developmental Synapse Length and Bouton Area Comparisons. **A**. Gardner-Altman estimation plot (Gardner and Altman, 1986) showing the raw synapse cross section length values for >p90 and p60-66 groups (left y-axis), as well as the mean difference (black dot near right y-axis), and the bootstrap 95% confidence interval (vertical error bar from black dot). Mann Whitney U test, p value = 1.5 × 10^−4^. **B**. Estimation plot showing raw excitatory bouton cross section areas for all developmental age groups examined. Plots marked as in A, except that mean differences between groups and confidence intervals are shown below each comparison. Mann Whitney U test, p value = 0.004 (p60-66 vs. >p90), and 2.8 × 10^−14^ (p21-28 vs. p46-47).

In 2D TEM images, spinule cross-sections (i.e. 2D profiles of embedded putative spinules) were present within excitatory synaptic boutons at every age examined. Spinule cross-sections within excitatory boutons displayed two lipid bilayers (outer bouton and inner spinule membranes) and were distinguished from other cellular organelles (see Methods). In measuring spinule areas within SBBs across postnatal development, we found that average spinule area remained relatively constant but showed a trend that mirrored the development of bouton areas from before eye-opening until after p90 (**Fig. 2C, Tables 1 & 3**). Moreover, on average spinules occupied ~6-11% of their encapsulating bouton’s area over development, and this relationship held steady until after p90 (**Table 1**). Yet despite this relatively stable spinule area to bouton area ratio, the SBB population at >p90 had larger bouton cross section areas than >p90 boutons without spinules (**Fig. 2-2, Table 3**). These data suggest that spinule engulfment increases SBB size over cortical boutons without spinules, and that a mechanism may exist to constrain spinule size as they invaginate into larger SBBs over development.

**Figure 2-2.**
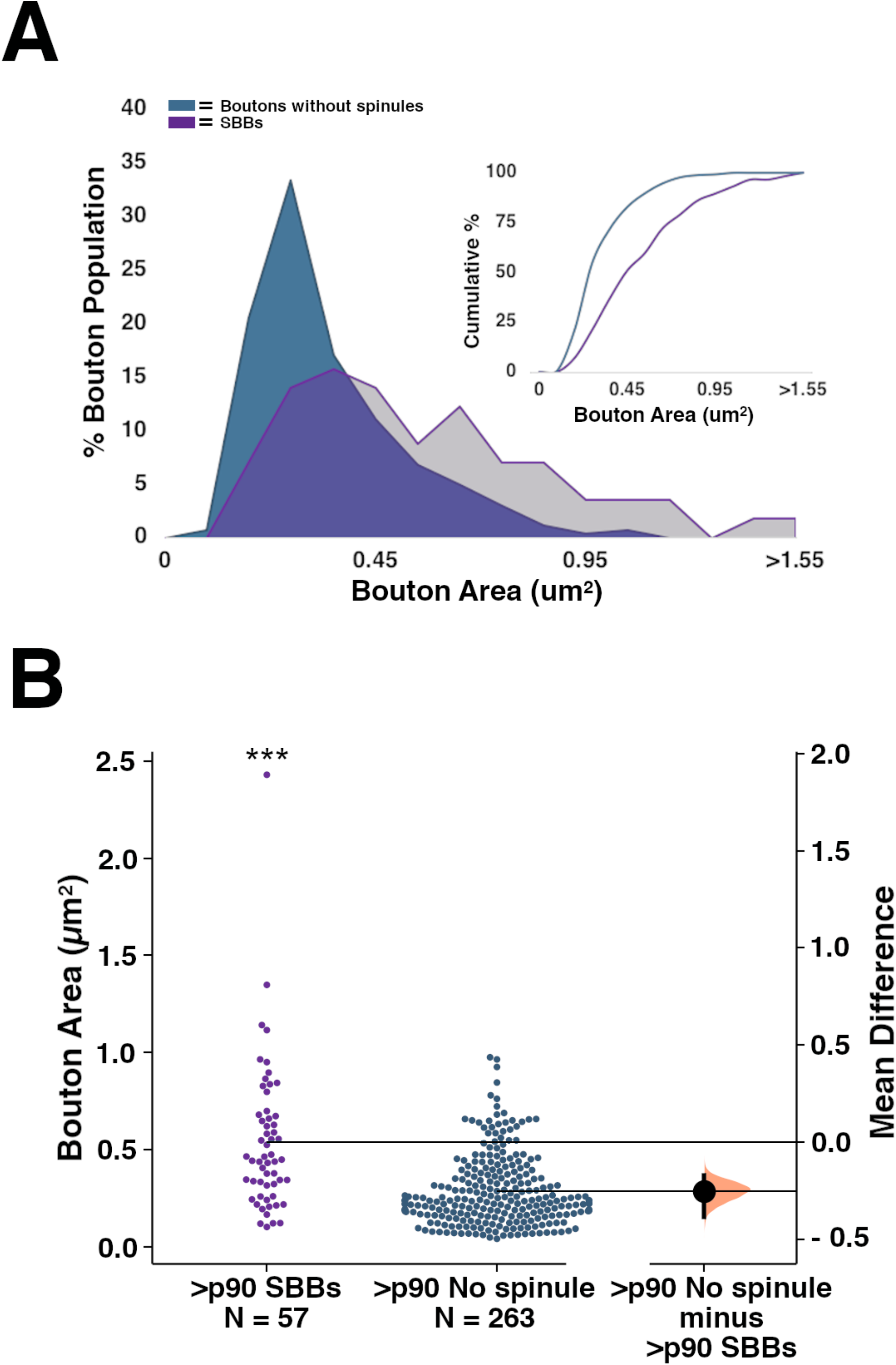
Spinule-Bearing Boutons (SBBs) are Larger than Boutons Without Spinules. **A**. Histogram showing the distribution for the percentage of >p90 cortical SBBs (purple) and >p90 presynaptic boutons without spinules (blue) with various sized cross section areas. Inset: Cumulative histogram of this same comparison. **B**. Estimation plot showing the raw data, mean difference (black dot), and bootstrapped 95% confidence interval (lines extending from dot) for cross section areas from >p90 SBBs and boutons without spinules. Mann Whitney U test, p = 2.7 × 10^−9^. *** = p < 0.001.

### Spinule Prevalence Across Development: 2D TEM Analyses

We hypothesized that if spinule engulfment by SBBs were driven by the onset of sensory activity or a heightened state of cortical plasticity, SBBs should increase in prevalence between eye-opening (~p30) and the height of Hebbian and homeostatic plasticity in V1 (~p46-47) (Issa et al., 1999; Yu et al., 2011). Furthermore, we predicted that if spinule engulfment by cortical boutons were correlated with the state of V1 plasticity, SBBs should be most prevalent around p46-47, and then should slowly wane in parallel with the expression of plasticity in ferret binocular V1 (Issa et al., 1999; Li et al., 2006). In contrast, if spinules play a role in stabilizing functionally and morphologically mature synapses, SBB prevalence should continue to increase as the expression of cortical plasticity wanes. We analyzed 910 excitatory presynaptic boutons across ferret postnatal developmental for the presence of spinules and found that the percentage of SBBs in L4 did not appreciably change from before the onset of visual experience until the canonical peak of plasticity in V1 (**Fig. 2D, Tables 1 & 3**).

In contrast, the percentage of SBBs trended toward increased levels between the height (~p46-47) and end of the canonical critical period in V1 (p60-66; p = 0.041, not significant), and the proportion of cortical boutons containing a spinule significantly increased between p60-66 and >p90 (68% increase; **Fig. 2D, Tables 1 & 3**). Thus, although deprivation-induced cortical plasticity within ferret V1 decreases by ~50% between p42 and p60, and deprivations after p100 fail to induce measurable ocular dominance plasticity (Issa et al., 1999), the percentage of SBBs within ferret V1 increase significantly over this same time frame. Taken together, these data suggest that spinule encapsulation by developing cortical SBBs is regulated by processes that parallel the physiological and morphological maturation and presynaptic stability of V1 synapses (Jones and Calverley, 1991; Durack and Katz, 1996; Erisir and Dreusicke, 2005; Cheetham and Fox, 2010; Smith et al., 2015; Qiao et al., 2016), and that SBB prevalence is not correlated with the onset of sensory activity or with the morphological malleability permitted during heightened states of neocortical plasticity.

### 3D FIBSEM Analyses of SBBs in V1

We took advantage of both 2D TEM and 3D focused ion beam scanning electron microscopy (FIBSEM) in a complementary approach to determine the time course for the developmental expression of SBBs in L4 of V1. Our 2D TEM analyses were designed to quantify a large number of excitatory boutons across multiple animals and ultrathin sections per age group in order to capture the potential variability in SBB prevalence and quantitative morphometry across individual animals and cortical depth. However, while they have higher lateral resolution, 2D TEM analyses are potentially biased toward omitting smaller objects (i.e. smaller boutons and spinules) (Colonnier and Beaulieu, 1985). Moreover, due to a larger physical section thickness (i.e. ~50-80 nm vs. 4 nm in FIBSEM), 2D TEM analyses are not suitable for quantifying small (e.g. 20-30 nm) perforations in PSDs, and analyses of single TEM images cannot be used to determine the neurite/glial source of spinules embedded within SBBs. Thus, in order to determine the origin of spinules within SBBs, the potential relationship between an invaginating spinule and perforations in a synaptic bouton’s PSD, and ascertain the accuracy of our 2D analyses of SBB prevalence, we prepared one tissue block from each developmental age group (i.e. p21, p46, p60, and >p90) for FIBSEM. We obtained 4 nm/pixel isotropic FIBSEM image stacks of L4 V1 from each animal, yielding aligned image volumes from p21 (15.1 × 14.1 × 2.8 μm), p46 (9.7 × 8.4 × 2.7 μm), p60 (24.2 × 16.2 × 2.4 μm), and >p90 (24.2 × 16.2 × 2.4 μm) brains. We evaluated every excitatory synapse we encountered (**Table 3**) within each image volume in 3D for the presence/absence of a spinule within the synaptic bouton, the origin of the invaginating spinule (if present), and the presence/absence of a perforated PSD. In addition, we reconstructed a total of 22 SBBs along with their invaginating spinules, postsynaptic targets, and PSDs (**Figs. 3 – 7**). With 4 nm isotropic pixel resolution we were able to follow every spinule back to its parent neurite, as well as determine the percentages of PSDs that contained perforations. Since perforations in the PSD of cortical synapses have been found to correlate with measures of synaptic plasticity (Calverley and Jones, 1990; Geinisman et al., 1993; Toni et al., 1999; Ganeshina et al., 2004), and potentially with the prevalence of SBBs (Jones et al., 1991; Spacek and Harris, 2004), we evaluated each synapse for the presence of perforated PSDs.

**Figure 3.**
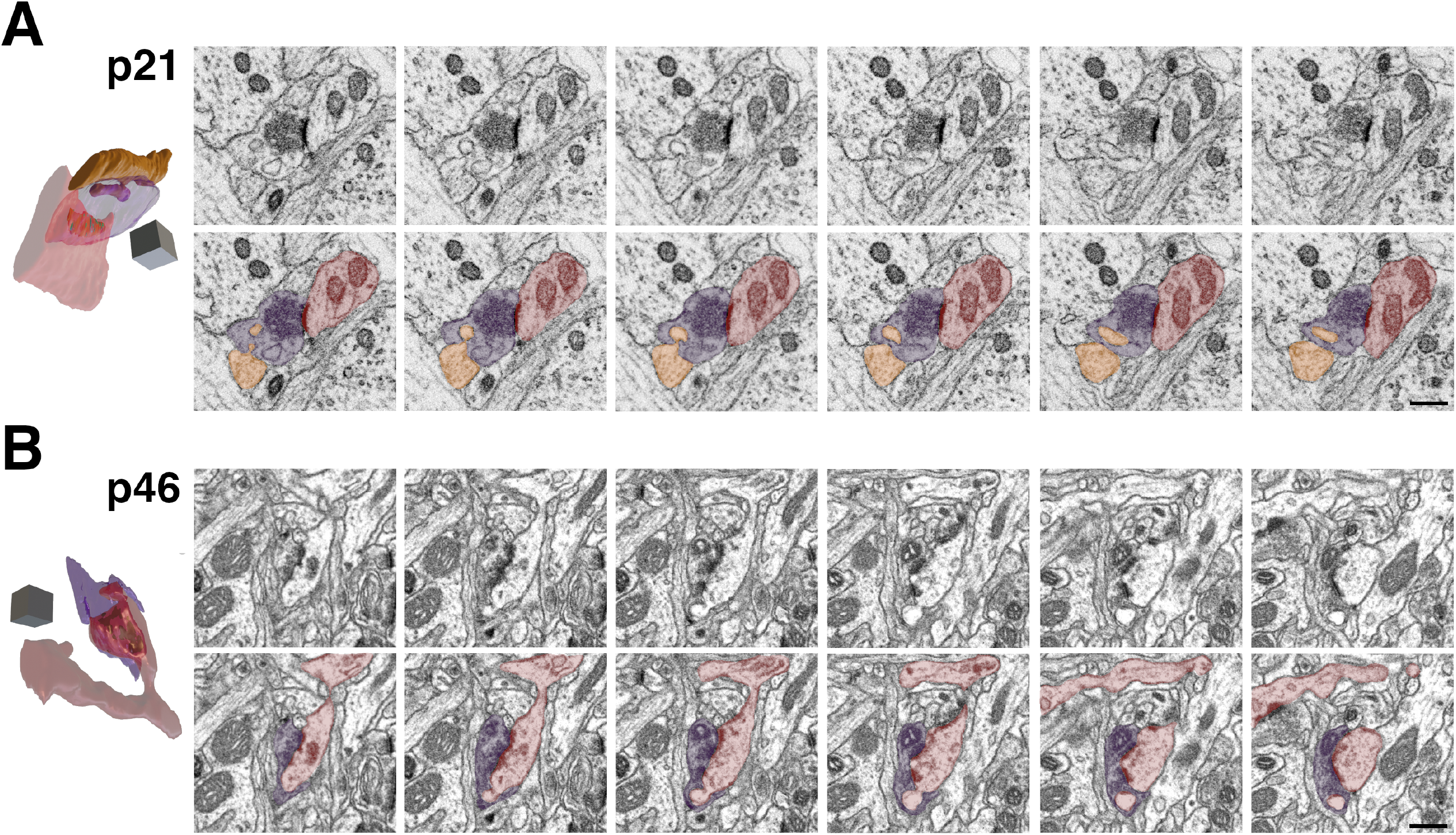
Focused Ion Beam Scanning Electron Microscopy (FIBSEM) Images of Spinule-Bearing Boutons (SBBs) in p21 and p46 V1. **A**. Adjacent FIBSEM images of a SBB in L4 of V1 from a p21 ferret. FIBSEM images are ~25 nm apart in z (depth) on average, picked to display the progression of spinule invagination and engulfment into the SBB. Top panel shows raw FIBSEM images, bottom panel is pseudo-colored to highlight SBB (purple), adjacent axon projecting a spinule (orange), and postsynaptic dendrite (red). At Left: Full reconstruction of this SBB showing transparent bouton with engulfed adjacent axon spinule and postsynaptic dendrite. and bouton spinules. Note the macular shaped postsynaptic density (PSD; green) formed between the SBB and the red dendrite. Identical SBB as shown in Fig. 5A & A_1_. **B**. Adjacent FIBSEM images of a SBB in L4 of V1 from a p46 ferret. Top and bottom panels arranged and colored as in A, showing a postsynaptic spine (red) projecting a spinule into its presynaptic SBB partner (purple). At Left: Full reconstruction of this p46 SBB showing postsynaptic spine spinule enveloped by its presynaptic bouton. Note the perforated complex shaped PSD (green). Identical SBB as shown in Fig. 5D & D_1_. Scale bars = 0.5 μm; 3D scale cubes = 0.5 μm^3^.

We encountered a range of neurites that projected spinules into L4 SBBs, including postsynaptic spines, adjacent (i.e. not synaptic with the SBB) spines, adjacent dendrites, adjacent axons/boutons, and glial processes, in line with previous reports from hippocampus (Sorra et al., 1998; Spacek and Harris, 2004). Moreover, spinules embedded within SBBs seemed to have a range of sizes and shapes, even within a single spinule origin category (e.g. postsynaptic spine spinules). Some spinules, particularly from select postsynaptic spines and adjacent axons, had thin initial invaginations into their SBB but then expanded into an anchor or hook-like shape (**Figs. 5A, 6B, 7D**). In addition, SBBs sometimes enveloped a portion of the head of their postsynaptic spine partners (**Fig. 6A, 7C, 7F**), a phenomenon previously described for TC boutons and their postsynaptic spines in mouse barrel cortex (Rodriguez-Moreno et al., 2018). Rarely, SBBs in older animals were observed enveloping spinules from multiple neurites (5.6 % and 1.9 % of SBBs for p60 and >p90, respectively; **Figs. 4A, 5E-F, 6C**). Thus, SBBs in L4 of V1 contain a diverse array of spinules from postsynaptic and adjacent neurites and glia, possessing a range of morphologies suggestive of distinct functions.

**Figure 4.**
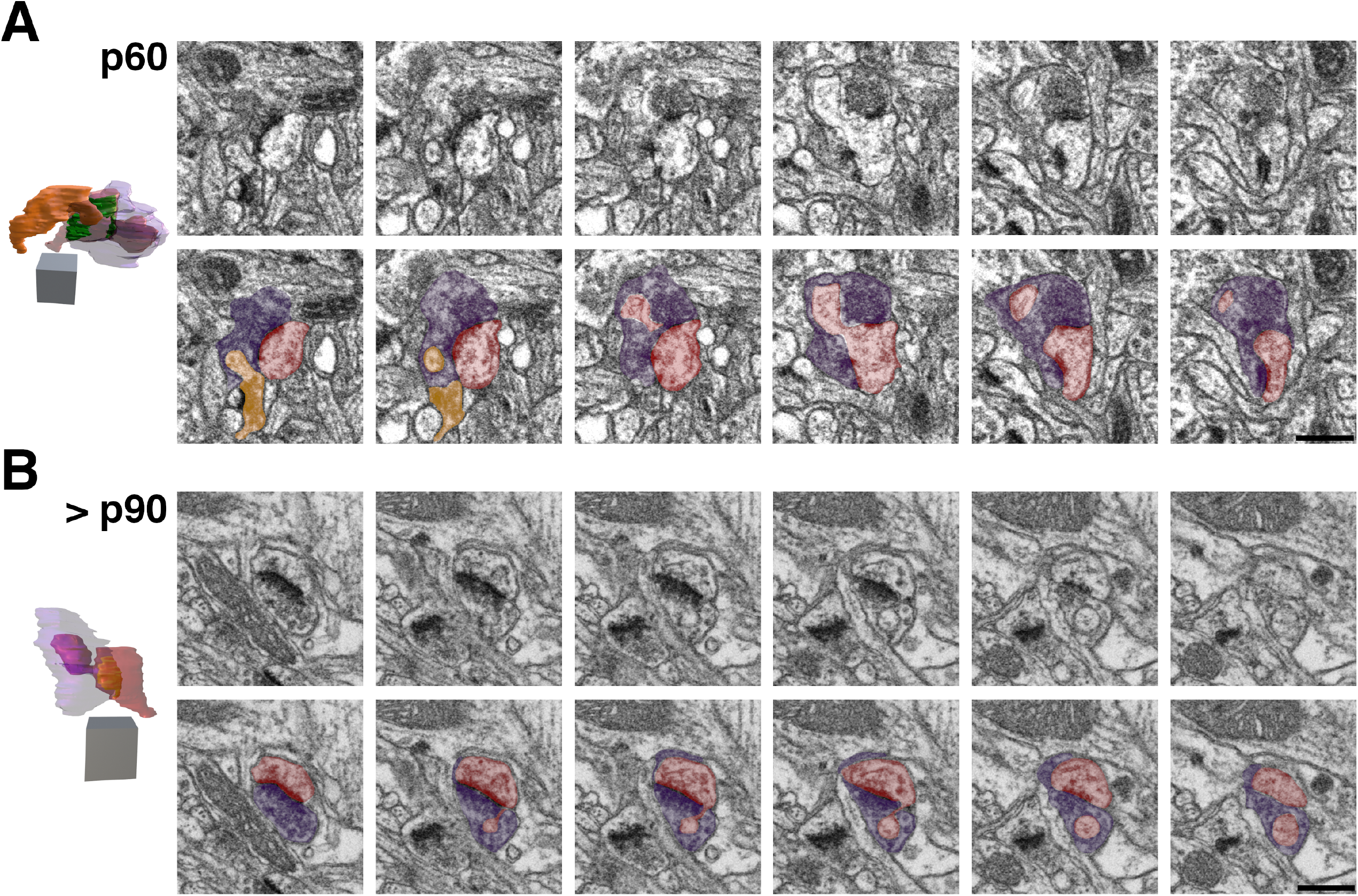
Focused Ion Beam Scanning Electron Microscopy (FIBSEM) Images of Spinule-Bearing Boutons (SBBs) in p60 and >p90 V1. **A**. Adjacent FIBSEM images of a SBB in L4 of V1 from a p60 ferret. FIBSEM images are ~25 nm apart in z (depth) on average, picked to display the progression of spinule invagination and engulfment into the SBB. Top panel shows raw FIBSEM images, bottom panel is pseudo-colored to highlight SBB (purple), adjacent axon/bouton (orange), and postsynaptic spine (red). Note the adjacent bouton (orange) with a synapse onto a spine that protrudes a spinule into this SBB at the bottom left of the first image in the series, and the spinule from the postsynaptic spine that invaginates into this SBB across the middle of its perforated postsynaptic density (PSD). At Left: Full reconstruction of this SBB showing transparent bouton (purple) with engulfed postsynaptic spine (red) and adjacent bouton (orange) spinules. Note the horseshoe-shaped perforated PSD. Identical SBB as shown in Fig. 6C & C_1_. **B**. Adjacent FIBSEM images of a SBB in L4 of V1 from a >p90 ferret. Top and bottom panels arranged and colored as in A. Note the postsynaptic spine (red) that sends its spinule into its SBB (purple) partner from the edge of the PSD. At Left: Full reconstruction of this SBB (purple), made transparent to show the engulfed anchor-like spinule from its postsynaptic spine partner. Macular-shaped PSD (green) appears yellow within spine. Identical SBB as shown in Fig. 7D & D_1_. 2D scalebars = 0.5 μm; 3D scale cubes = 0.5 μm^3^.

**Figure 5.**
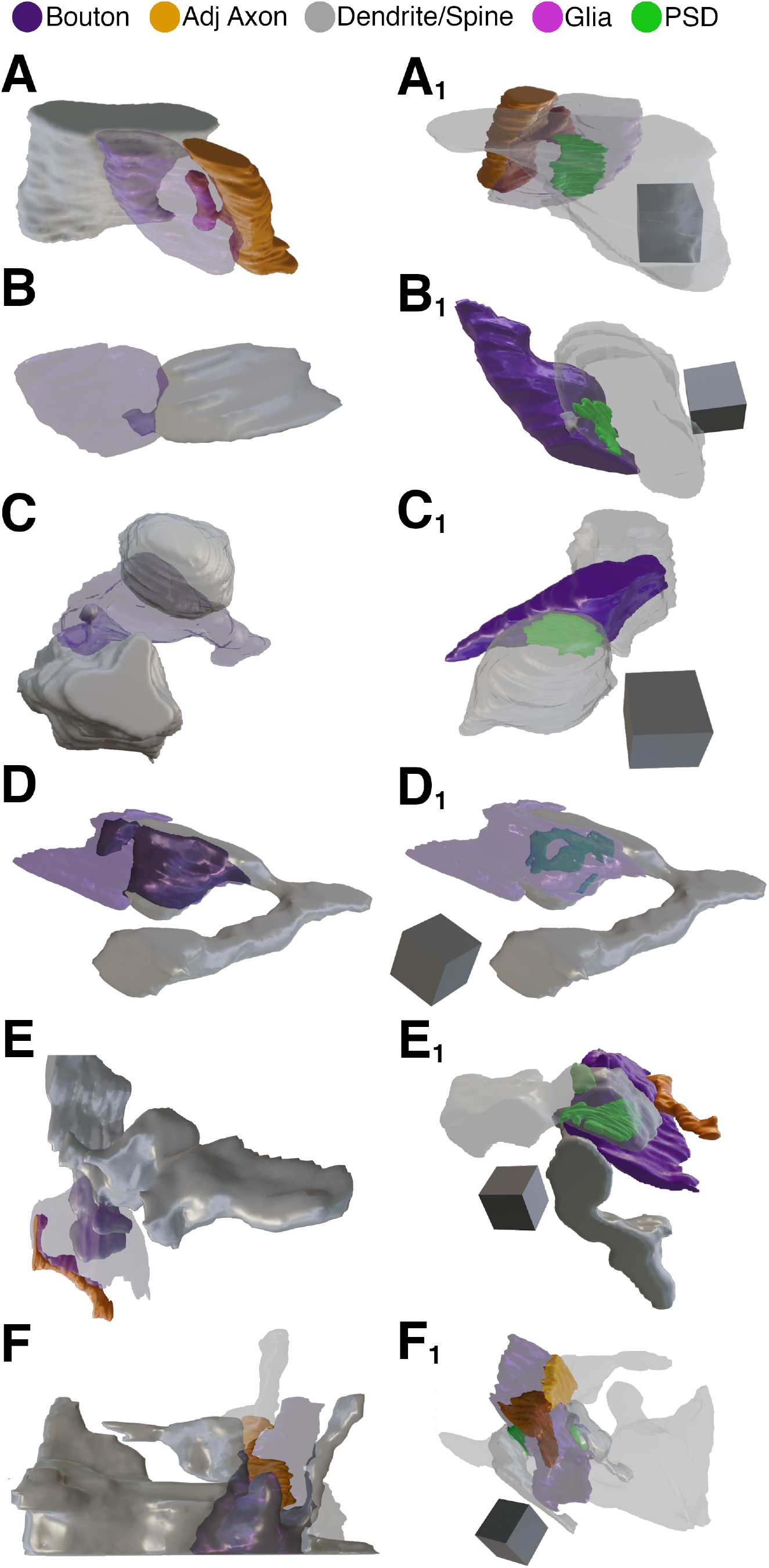
3D Reconstructions of p21 and p46 Spinule-bearing Boutons (SBBs). Focused ion beam serial electron microscopy 3D reconstructions of SBBs (purple) and their spinules from axon/boutons (yellow), and spines/dendrites (gray). **A – C**. 3D reconstructions from a p21 ferret showing spinules from an adjacent axon (A and A_1_), postsynaptic dendrite (B and B_1_), and adjacent dendrite (C and C_1_) projecting into L4 SBBs. Note that in B, the spinule emerges from the edge of the postsynaptic density (PSD). Reconstruction shown in A is the identical SBB shown in Fig. 3A. **D - F**. 3D reconstructions from a p46 ferret showing hook-like spinule from a postsynaptic spine (D and D_1_), and spinules from adjacent axons and adjacent dendrites (E, E_1_, F, and F_1_) projecting into L4 SBBs. Note the complex perforated PSD in D1, the identical SBB shown in Fig. 3B. In F and F_1_, an SBB with synapses (green) onto two postsynaptic spines receives a large adjacent dendrite spinule (F) and a large adjacent axon spinule (F_1_). **A_1_ – F_1_**. Identical SBBs as shown to the left in A-F, but with transparent postsynaptic neurites and/or SBBs to highlight the morphology and locations of the postsynaptic densities (green) or spinules. 3D scale cubes = 0.5 μm^3^.

**Figure 6.**
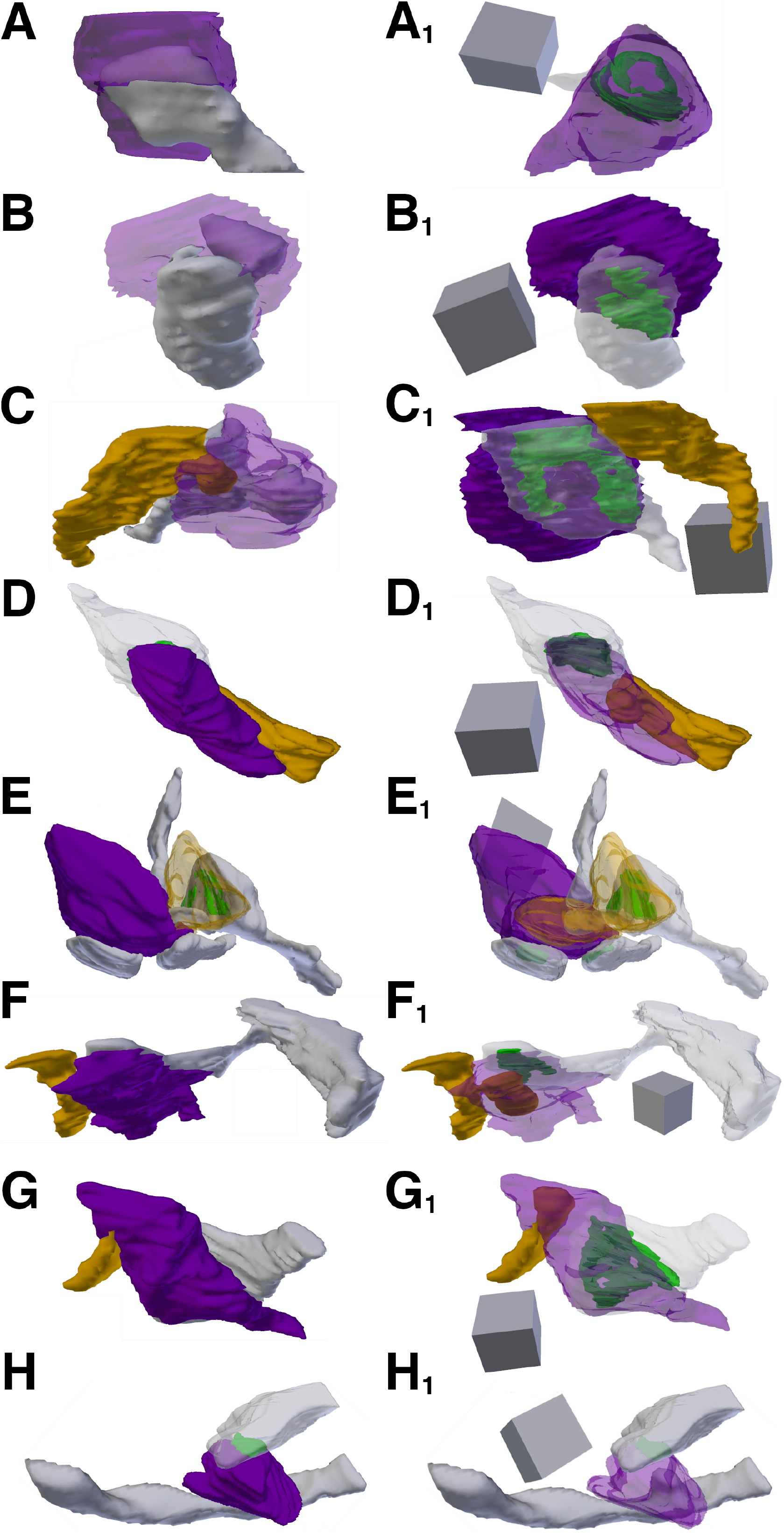
3D Reconstructions of p60 Spinule-bearing Boutons (SBBs). Focused ion beam serial electron microscopy 3D reconstructions of SBBs (purple) and their spinules from axon/boutons (yellow), and spines/dendrites (gray). **A – C**. Reconstructions of a postsynaptic spine head (A) and postsynaptic spine anchor-like spinules (B-C) projecting into L4 p60 SBBs. Note that in C, a second spinule from a synaptic bouton projects into the ‘upper’ portion of the bouton (same bouton shown in Fig. 4A). **C – G**. Reconstructions showing axons/bouton spinules of various sizes engulfed by p60 SBBs. Note that in E, a presynaptic bouton (yellow) receives a spinule from its postsynaptic spine partner and a portion of this bouton with its engulfed spine protrude into a large adjacent SBB (purple) with synapses (green) onto three postsynaptic spines. **H**. A SBB with a synapse onto a postsynaptic dendrite (top) receives a spinule from an adjacent dendrite (bottom). **A_1_ – H_1_**. Identical SBBs as shown to the left in A-H, but with transparent postsynaptic neurites and/or SBBs to highlight the morphology and locations of the postsynaptic densities (green) or spinules. 3D scale cubes = 0.5 μm^3^.

**Figure 7.**
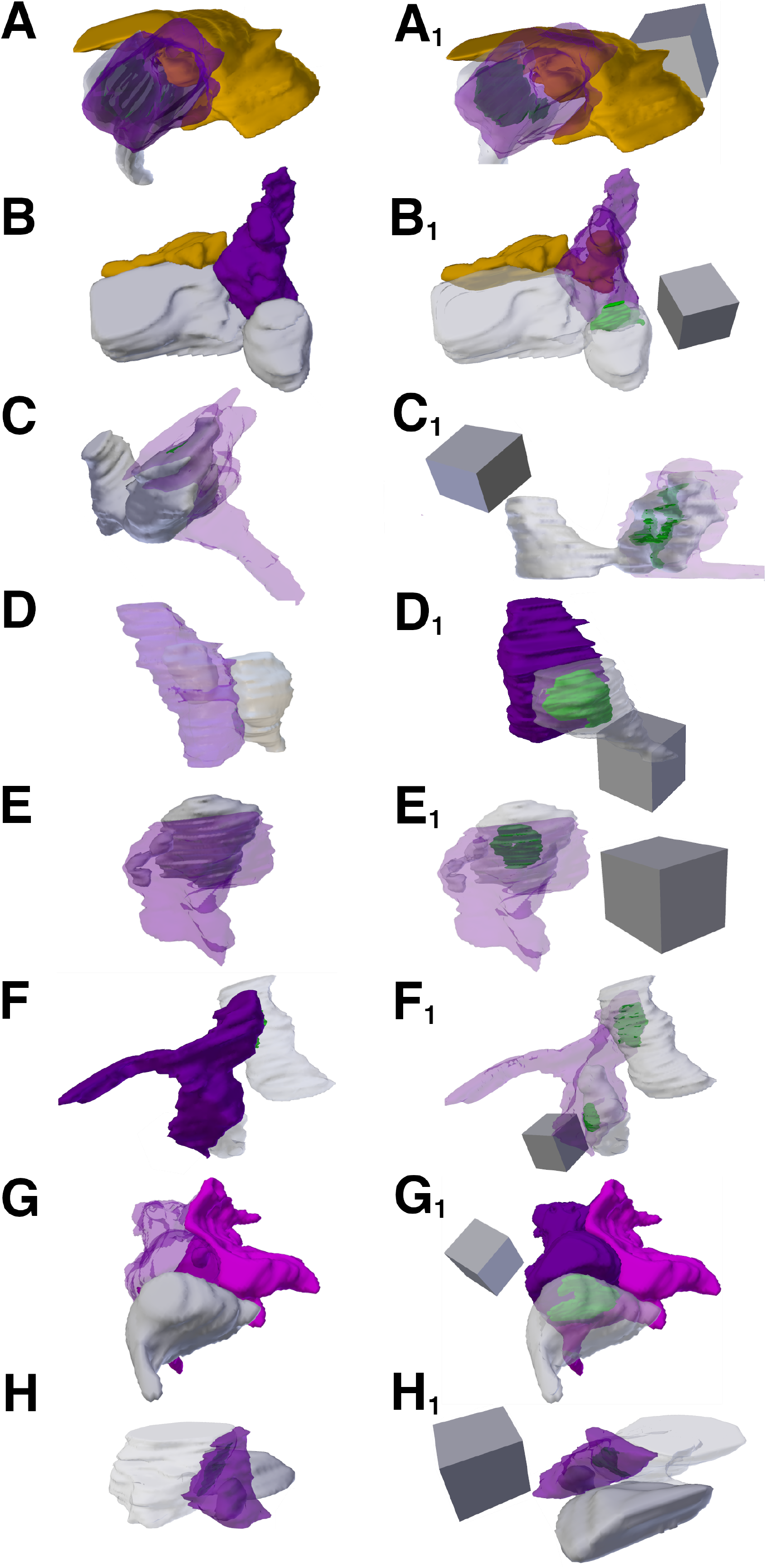
3D Reconstructions of >p90 Spinule-Bearing Boutons (SBBs). Focused ion beam serial electron microscopy 3D reconstructions of SBBs (purple) and their spinules from axon/boutons (yellow), spines/dendrites (gray), and glia (pink) from L4 of V1 of a >p90 ferret. **A – B**. Reconstructions showing adjacent axons/bouton spinules engulfed by SBBs with synapses onto postsynaptic spines. **C – F**. Reconstructions of postsynaptic spine heads (C and F), and anchor-like spinules (D and E) projecting into their SBB partners. Note the horseshoe-shaped perforated postsynaptic density (green) in C_1_, and that the SBB shown in F and F_1_ envelops approximately two-thirds of its postsynaptic spine partner. Reconstruction shown in D is the identical SBB shown in Fig. 4B. **G**. A SBB with a synapse onto a postsynaptic spine receives a spinule from an adjacent glia. **H**. A SBB with a synapse onto a postsynaptic dendrite engulfs a spinule from an adjacent dendrite. **A_1_ – H_1_**. Identical SBBs as shown to the left in A-H, but with transparent postsynaptic neurites and/or SBBs to highlight the morphology and locations of the postsynaptic densities (green) or spinules. 3D scale cubes = 0.5 μm^3^.

### Spinule Prevalence Across Development: 3D FIBSEM Analyses

Our quantitative 3D FIBSEM analyses of 676 excitatory boutons revealed that the percentage of SBBs within the synaptic bouton population of L4 remains at ~6.5% from before eye opening (p21) until the height of the critical period for plasticity (p46) (**Fig. 8A**). Yet by p60, 14.5% of excitatory boutons contain spinules (a 127% from p46 levels), and by >p90 spinules are embedded within 24.2% of excitatory boutons (67% increase from p60 levels; **Fig. 8A, Tables 2 & 3**). These data mirror the trends we observed in our 2D TEM analyses of spinule prevalence across these same developmental time points (**Fig. 8A**). Together, these analyses demonstrate that the rate and/or maintenance of spinule engulfment by excitatory boutons in L4 of V1 is most likely regulated by mechanisms that parallel the waning of cortical plasticity, suggesting spinules act to stabilize functionally and/or morphologically mature neocortical synapses.

**Table 2.** Percentages of SBBs, Perforated PSDs, Spinule Origins, and Sampled Structures for FIBSEM Analyses. Total sampled boutons and synapses for each age are listed under “Total Boutons” and “Total Synapses” columns for FIBSEM analyses at each age examined. Percentages of spinule-bearing boutons (SBBs) and perforated postsynaptic densities (PSDs) in FIBSEM analyses are listed in their respective columns for each age examined. Percentages of SBBs containing at least one spinule from a neurite/glial source are listed in their respective columns for each age examined. PS = postsynaptic; Adj = adjacent (i.e. not synaptic); Unknown = spinules whose origin could not be determined (see Methods for details). Since some SBBs contained spinules from multiple sources, percentages may not sum to 100 %.

| Age | Total Boutons | Total Synapses | % SBBs | % Perforated PSDs | % PS Spine | % Adj Axon | % Adj Spine | % Adj Dendrite | % Adj Glia | % Unknown |
| --- | --- | --- | --- | --- | --- | --- | --- | --- | --- | --- |
| >p90 | 223 | 237 | 24.2 | 30.5 | 37.0 | 35.2 | 7.4 | 14.8 | 5.6 | 3.7 |
| p60 | 235 | 256 | 14.5 | 16.8 | 14.7 | 55.9 | 8.8 | 17.7 | 0.0 | 0.0 |
| p46 | 157 | 163 | 6.4 | 16.6 | 10.0 | 50.0 | 0.0 | 40.0 | 0.0 | 0.0 |
| p21 | 61 | 62 | 6.6 | 16.1 | 25.0 | 25.0 | 0.0 | 25.0 | 0.0 | 25.0 |

**Table 3.** Statistical Comparisons for 2D Transmission Electron Microscopy (TEM) and 3D Focused Ion Beam Scanning Electron Microscopy (FIBSEM) Analyses. Statistical tests used for 3D analyses are listed, grouped by each quantified morphology category (left most column) and age/species comparison (“Comparison” column). Percent change between means are calculated as: 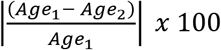, where *Age*_*n*_ = Mean of age/species group. Power is the post-hoc two-tailed achieved statistical power for each comparison, given the two proportions, sample sizes (n), with ∞ error probability = 0.05. * = comparisons where and p < 0.05 or Bonferroni corrected value. Sample sizes for ‘Spinule Origins’ comparisons reflect the total number of spinule-bearing boutons (SBBs) within each group. PSDs = Postsynaptic densities. 2D = TEM analyses.

| Comparison | Test | n (animals, total sampled structures) | % Change | Power (post-hoc, two-tailed) | p value |
| --- | --- | --- | --- | --- | --- |
| **Synapse Length** |  |  |  |  |  |
| All Ages | ANOVA, one-way |  |  |  | 6.7 x 10<sup>-7</sup>* |
| >p90 vs. p60-66 | Bonferroni post hoc t Test, two-tailed | >p90 (3, 323); p60-66 (3, 266) | 16% | 0.96 | 2.1 x 10<sup>-4</sup>* |
| p60-66 vs. p46-47 | Bonferroni post hoc t Test, two-tailed | p60-66 (3, 266); p46-47 (2, 196) | 5% | 0.14 | 0.327 |
| p46-47 vs. p21-28 | Bonferroni post hoc t Test, two-tailed | p46-47 (2, 196); p21-28 (2, 152) | 2% | 0.07 | 0.669 |
| **Bouton Area** |  |  |  |  |  |
| All Ages | ANOVA, one-way |  |  |  | 6.0 x 10<sup>-13</sup>* |
| >p90 vs. p60-66 | Bonferroni post hoc t Test, two-tailed | >p90 (3, 319); p60-66 (3, 254) | 20% | 0.82 | 0.005* |
| p60-66 vs. p46-47 | Bonferroni post hoc t Test, two-tailed | p60-66 (3, 254); p46-47 (2, 196) | 39% | 0.98 | 6.1 x 10<sup>-5</sup>* |
| p46-47 vs. p21-28 | Bonferroni post hoc t Test, two-tailed | p46-47 (2, 196); p21-28 (2, 141) | 93% | 0.99 | 1.34 x 10<sup>-11</sup>* |
| **Spinule Area** |  |  |  |  |  |
| All Ages | ANOVA, one-way |  |  |  | 0.671 |
| **Spinule-Bearing Bouton Area** |  |  |  |  |  |
| >p90 SBBs vs. >p90 No Spinules | Mann-Whitney U test | > p90 SBBs (3, 57); >p90 NoSpin (3, 266) | 88% | 0.99 | 2.69 x 10<sup>-9</sup>* |
| **% Spinule-Bearing Boutons** |  |  |  |  |  |
| **2D** |  |  |  |  |  |
| All Ages (4 x 2 table) | X<sub>2</sub> Test, two-tailed, Bonferroni corrected |  |  |  | 5.3 x 10<sup>-5</sup>* |
| >p90 vs. p60-66 | X<sub>2</sub> Test, two-tailed, Bonferroni corrected | >p90 (3, 323); p60-66 (3, 254) | 68% | 0.73 | 0.009* |
| p60-66 vs. p46-47 | X<sub>2</sub> Test, two-tailed, Bonferroni corrected | p60-66 (3, 254); p46-47 (2, 196) | 92% | 0.52 | 0.041 |
| p46-47 vs. p21 | X<sub>2</sub> Test, two-tailed, Bonferroni corrected | p46-47 (2, 196); p21-28 (2, 141) | 8% | 0.04 | 0.863 |
| **FIBSEM** |  |  |  |  |  |
| All Ages (4 x 2 table) | X<sub>2</sub> Test, two-tailed, Bonferroni corrected |  |  |  | < 1.0 x 10<sup>-5</sup>* |
| >p90 vs. p60 | X<sub>2</sub> Test, two-tailed, Bonferroni corrected | >p90 (1, 223); p60 (1, 235) | 67% | 0.81 | 0.008* |
| p60 vs. p46 | X<sub>2</sub> Test, two-tailed, Bonferroni corrected | p60 (1, 235); p46 (1, 157) | 127% | 0.70 | 0.014* |
| p46 vs. p21 | X<sub>2</sub> Test, two-tailed, Bonferroni corrected | p46 (1, 157); p21 (1, 61) | -3% | 0.03 | 0.959 |
| **% Perforated PSDs** |  |  |  |  |  |
| **FIBSEM** |  |  |  |  |  |
| All Ages (4 x 2 table) | X<sub>2</sub> Test, two-tailed, Bonferroni corrected |  |  |  | 0.002* |
| >p90 vs. p60 | X<sub>2</sub> Test, two-tailed, Bonferroni corrected | >p90 (1, 237); p60 (1, 256) | 81% | 0.94 | 7.0 x 10<sup>-4</sup>* |
| p60 vs. p46 | X<sub>2</sub> Test, two-tailed, Bonferroni corrected | p60 (1, 256); p46 (1, 163) | 2% | 0.04 | 0.882 |
| p46 vs. p21 | X<sub>2</sub> Test, two-tailed, Bonferroni corrected | p46 (1, 163); p21 (1, 62) | 3% | 0.04 | 0.937 |
| **Spinule Origins: >p90 vs. p60-66** |  |  |  |  |  |
| % Postsynaptic Spine (>p90 vs. p60) | X<sub>2</sub> Test, two-tailed, Bonferroni corrected | >p90 (1, 54); p60 (1, 34) | 152% | 0.63 | 0.024* |
| % Adjacent Axon (>p90 vs. p60) | X<sub>2</sub> Test, two-tailed, Bonferroni corrected | >p90 (1, 54); p60 (1, 34) | -37% | 0.48 | 0.030* |
| % Adjacent Dendrite (>p90 vs. p60) | X<sub>2</sub> Test, two-tailed, Bonferroni corrected | >p90 (1, 54); p60 (1, 34) | -16% | 0.06 | 0.724 |
| % Adjacent Spine (>p90 vs. p60) | X<sub>2</sub> Test, two-tailed, Bonferroni corrected | >p90 (1, 54); p60 (1, 34) | -16% | 0.05 | 0.271 |
| % Adjacent Glia (>p90 vs. p60) | X<sub>2</sub> Test, two-tailed, Bonferroni corrected | >p90 (1, 54); p60 (1, 34) | n/a | 0.24 | 0.161 |

**Figure 8.**
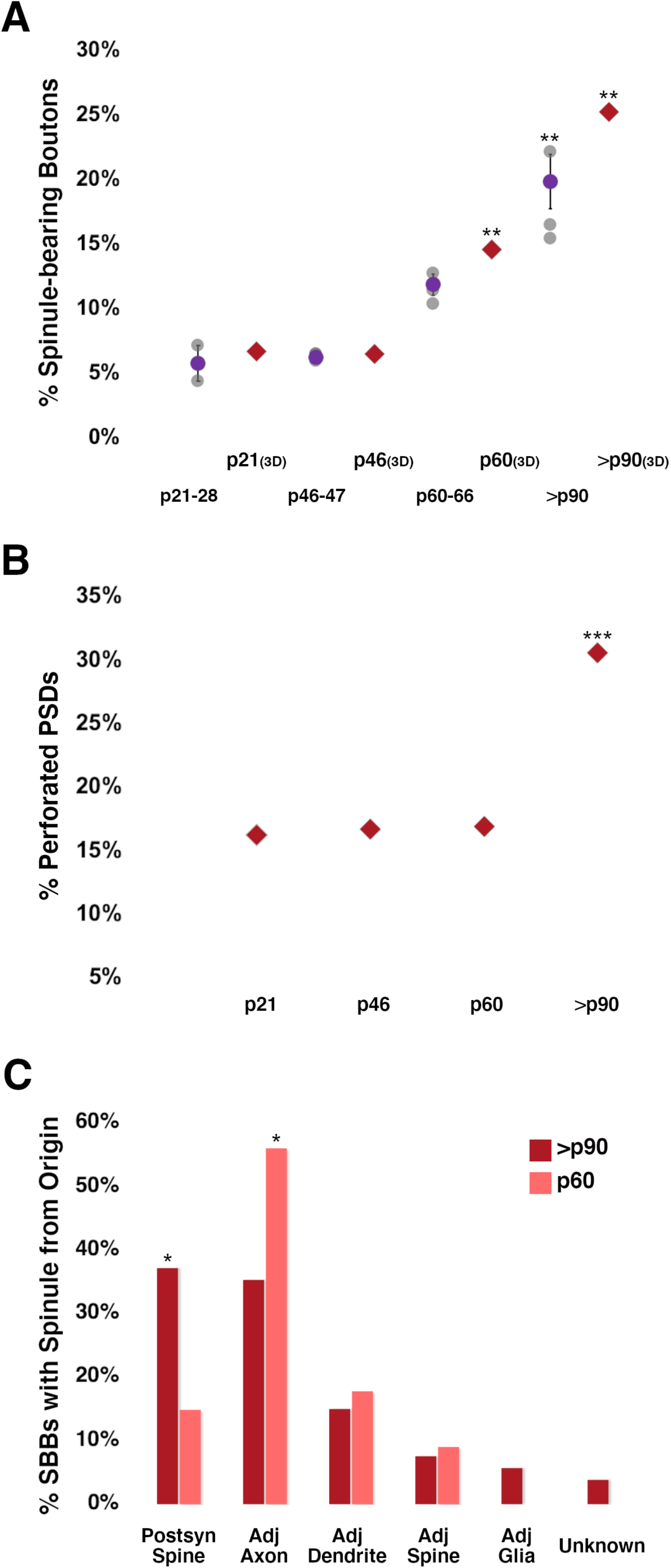
Spinule-Bearing Boutons Progressively Engulf Spinules from the End of the Canonical Critical Period into Late Adolescence. **A**. Percentages of excitatory spinule-bearing boutons (SBBs) across the developmental ages examined in 2D TEM and FIBSEM (3D) images. 3D FIBSEM data at p21, p46, p60, and >p90 are represented by red diamonds, 2D data as shown in Fig. 2D. **B**. FIBSEM developmental analysis showing the percentages of excitatory presynaptic boutons with perforated postsynaptic densities (PSDs). **C**. FIBSEM analysis showing the percentages of p60 and >p90 SBBs containing at least one spinule from a defined neurite or glial origin. For example, ~56% of SBBs at p60 contain at least one spinule from an adjacent axon. * = p < 0.05; ** = p < 0.015; *** = p < 0.001.

### Perforated PSDs and SBB Prevalence

As the prevalence of separations/perforations in PSDs have been suggested to correlate with levels of plasticity and spinule prevalence (Jones et al., 1991; Spacek and Harris, 2004), we sought to determine the developmental trajectory for the appearance of perforations in PSDs at excitatory synapses in our data. We reasoned that if spinules and perforated PSDs were both linked to heightened levels of plasticity and/or if spinules were responsible for producing perforations in PSDs as they invaginate into boutons, that spinule prevalence and levels of perforated PSDs would increase in parallel. However, we found that levels of perforated PSDs at excitatory synapses remained stable between p21 – p60 (~16-17%), followed by a significant increase to 30.5% by >p90 (**Fig. 8B, Tables 2 & 3**). Furthermore, the percentage of perforated PSDs at synapses formed by SBBs did not appreciably change across ages (50%, 40%, 35.3%, and 48.2% for p21, p46, p60, and >p90 respectively, Chi-Square test, 4×2 contingency table, p = 0.56), though our power to detect changes in perforated synapses at p21 and p46 SBBs is low given the small percentages of SBBs at these ages. Nevertheless, there does not appear to be a clear relationship between total levels of SBBs and perforated PSDs in V1 across the ages examined.

### Origins of Spinules within SBBs

Our FIBSEM analyses also allowed us to track the parent structure within the neuropil for every spinule embedded within our stacks of developing SBBs in L4 of V1. We hypothesized that spinules from postsynaptic spines, particularly larger anchor-like spinules (**e.g. Fig. 6B & 7D**) embedded within SBBs might encourage the maintenance and stability of synaptic connections. In contrast, spinules from non-synaptic adjacent neurites (e.g. axon/boutons, dendrites) might enable novel forms of communication or neuronal circuit remodeling (Spacek and Harris, 2004; Petralia et al., 2018). We focused on our analysis of spinule origins on p60 and >p90 ages where we had sufficient SBBs (see **Table 3**) to attain reasonable statistical power when comparing subsets of SBBs containing spinules from unique neurite/glial sources. We found that at both of these ages, SBBs primarily contained spinules from postsynaptic spines and adjacent axons/boutons, with a minority from other sources (**Fig. 8C; Table 2**). However, at p60 a majority of SBBs (56%) had spinules emanating from adjacent axons, and only ~15% of SBBs contained spinules from their postsynaptic spine partners. Yet by >p90, a significantly smaller percentage of SBBs contained spinules from adjacent axons (~35%), while a significantly larger percentage of SBBs contained spinules from their postsynaptic spine partners (37%). These data indicate that SBBs in L4 of V1 increasingly engulf spinules from postsynaptic spines as they mature, potentially acting to facilitate the stability of these connections. Whereas, SBBs decrease their preference for spinules from adjacent axons by >p90, potentially signaling the decreasing importance of these spinules for enabling extrasynaptic communication and/or circuit remodeling at more mature synapses.

## Discussion

Despite decades of discussion within the TEM literature, the developmental prevalence and origins of spinules within cortical boutons has remained obscure. In this study we performed 2D TEM and 3D FIBSEM analyses to determine the proportion of excitatory cortical boutons that envelop spinules across ferret postnatal development in V1 and quantified the percentages of spinules embedded within neocortical boutons that arise from unique neurite/glial sources. We found that: (1) the proportion of SBBs within the L4 excitatory bouton population increases as physiological and morphological plasticity wanes, suggesting spinules play a role in stabilizing mature cortical synapses, (2) that SBBs preferentially engulf spinules from postsynaptic spines and adjacent boutons/axons by p60 and into late adolescence, and that (3) nearly one-quarter of the excitatory synaptic bouton population in late adolescent ferret V1 contain spinules.

### 2D TEM vs. 3D FIBSEM Considerations

In order to address inter-animal variability and cortical depth in our developmental analyses, we characterized cortical bouton developmental morphology and the percentages of cortical SBBs (n = 10 animals, 910 synaptic boutons) using 2D TEM, and then used some of the same exact tissue to perform new 3D FIBSEM analyses (n = 4 animals, 676 synaptic boutons). While we took great care to only sample each synapse once and to analyze each synaptic profile we encountered, there are limitations associated with 2D TEM analyses. Since small synapses may be undersampled in a 2D analysis (Colonnier and Beaulieu, 1985), it is possible that we failed to include some smaller synapses in our 2D analyses. While we cannot completely rule out this possibility, our 3D FIBSEM data provide strong confirmatory support for both the trends and relative differences in SBB prevalence across the ages we examined (**Fig. 8A, Table 3**). In addition, our 2D analyses were able to demonstrate the variability in SBB prevalence present across animals (e.g. 15 –22% across 3 animals at >p90) and suggest that SBBs are larger than boutons without spinules (**Fig. 2-2**), a testable prediction for future FIBSEM experiments.

Furthermore, in order to exclude bouton organelles (e.g. endoplasmic reticula, lysosomes, synaptic vesicles) from our SBB analyses within 2D ~60 nm thick TEM sections, we adopted conservative criteria (see Methods) to positively identify membrane bound structures as spinules within 2D bouton cross sections. Accordingly, it is conceivable that our 2D TEM analyses failed to include some small spinules across developmental ages due to their small size and/or our stringent quantitative criteria. Indeed, we found slightly higher SBB prevalence at p60 and >p90 in our FIBSEM analyses (14.5% and 24.2% SBBs, p60 and >p90 respectively) versus our 2D TEM analyses (11 ± 0.8% and 18 ± 2%, Mean ± S.E.M, p60 and >p90 respectively) that are likely attributable to small spinules that were missed in our 2D analyses.

### Relationship of Perforated Synapses to Synaptic Spinules

Increases in the proportion of PSDs displaying perforations and ‘horseshoe’ morphology are associated with increases in cortical activity (Calverley and Jones, 1990; Geinisman et al., 1993; Toni et al., 1999; Harris et al., 2003), and can occur concomitant with increased spinule emergence from dendritic spines (Sorra et al., 1998; Toni et al., 1999; Spacek and Harris, 2004). Accordingly, one hypothesis for the functional significance of perforated PSDs might be the necessity of PSDs to perforate to allow for spinule growth into a bouton. However, similar to our observations in L4 of V1, a majority of spinules from hippocampal spines seem to emerge from the edge or distant from the PSD rather than from the middle of the PSD (Harris and Stevens, 1989; Spacek and Harris, 2004), suggesting that spinule emergence is not responsible for perforating most PSDs. Similarly, we found that increases in SBB prevalence preceded the developmental increase in perforated PSDs (**Fig. 8A – B**), and that only 35 – 50% of SBBs in V1 had synaptic partners with perforated PSDs, arguing against a causal relationship between spinule protrusions and perforated PSDs. Furthermore, our data are in line with multiple reports from rat neocortex where spinules have only been reported to appear within ~19% (Bozhilova-Pastirova and Ovtscharoff, 1999, rat S1), ~17% (Spacek, 1985, rat V1), and 13-37% (Jones and Calverley, 1991, rat parietal cortex across lifespan) of synapses with perforated PSDs. Hence, while our data demonstrate that spinules are increasingly engulfed by neocortical excitatory synaptic boutons over development, and the percentage of neocortical PSDs displaying perforations seems to increase as an animal matures (Geinisman et al., 1987; Calverley and Jones, 1990; Jones and Calverley, 1991; Dufour et al., 2016), it seems unlikely that spinule emergence or engulfment by SBBs are causally related to perforations in PSDs, at least in ferret V1. However, a recent live-imaging study of cortical neurons in culture found that dendritic spines protruding stable ‘long-lived’ spinules emanated farther away from their PSDs than short live spinules, but that long-lived spinules were more frequently associated with perforated PSDs (78%) than were spines projecting short-lived spinules (23%) (Zaccard et al., 2020). Thus, whether spinules directly influence the dynamic partitioning of PSDs, or whether this relationship holds true in vivo remain open questions.

### Nearly One-Quarter of Excitatory Cortical Boutons Contain Spinules

We found that nearly one-quarter (24.2%) of excitatory boutons in L4 of late adolescent (>p90) ferret V1 contain spinules. To our knowledge, there have only been two other published reports that investigated the proportion of excitatory bouton populations in vivo that contain spinules. Erisir & Dreusicke, (2005) found that in L4 of ferret V1, 10% of anterogradely-labeled thalamocortical (TC) boutons at p35-p49, and 28% of labeled TC boutons in adult contained spinules. Since, similar to our measurements of SBB boutons, this study found that labeled TC boutons are larger than their unlabeled counterparts, it is tempting to speculate that the SBBs that we sampled may be largely of TC origin. Indeed, a FIBSEM and serial TEM study of anterogradely labeled TC boutons in p60-65 mouse barrel cortex found that 40% of TC axospinous synapses had spinules or spine heads protruding into TC boutons (Rodriguez-Moreno et al., 2018). However, since spinules were not a focus of either of these TC studies, the variety, proportions, and specificity of spinule types for TC boutons versus non-TC intracortical boutons remain unclear. Moreover, L4 TC boutons only represent ~5-20% of the total excitatory bouton population in L4 of mammalian V1 (Ahmed et al., 1994; Peters et al., 1994; Latawiec et al., 2000; da Costa and Martin, 2009; Bopp et al., 2017; Garcia-Marin et al., 2019). Hence, given that 24.2% of excitatory boutons at >p90 were SBBs, it seems likely that at least a portion of the SBBs we encountered were intracortical excitatory boutons.

Examining SBBs within the TC population (e.g. VGluT2-containing) and excitatory intracortical bouton population (e.g. VGluT1-containing) are logical next steps in determining whether spinules from specific origins are preferentially engulfed by these functionally disparate classes of boutons. Yet, regardless of whether spinules are engulfed by specific bouton subtypes, our data suggest that hundreds to thousands of synapses on individual L4 cortical neurons (DeFelipe et al., 1999) contain SBBs. As such, spinules are poised to play prominent roles in facilitating synaptic stability (i.e. in SBBs physically connected to their postsynaptic spines), and potentially augmenting neuronal communication through transendocytosis (Spacek and Harris, 2004) and/or ephaptic coupling (Wagner and Djamgoz, 1993). Thus, an understanding of how spinules effect synapse function and/or stability will significantly impact our insight into neuronal input-output relationships.

### Spinule Origins and Potential Functions

To our knowledge, these data represent the first investigation into the origins of spinules within neocortical presynaptic boutons. A pioneering study into the 3D structure of spinules within CA1 hippocampus found that out of 254 identified spinules, a majority emerge from dendritic spines (~86%), and a minority emerge from axons (~12%) and dendritic shafts (~1%) (Spacek and Harris, 2004). Furthermore, these hippocampal spinules are engulfed by a majority of presynaptic boutons (~90%). The relative relationships between the proportions of spinule-projecting objects in hippocampus are in rough agreement with our data demonstrating that cortical boutons in L4 of late adolescent ferret engulf a majority of dendritic spines (44%: 37% postsynaptic spines and ~7% adjacent non-synaptic spines) and boutons/axons (35%). However, if SBBs comprise the majority of spinule-engulfing structures in primary sensory cortex as they seem to do in the hippocampus, there may be interesting systematic differences in the proportions of spinule projecting objects and/or SBB preference for specific spinule types between the hippocampus and neocortex. For example, axons/bouton spinules appear to be overrepresented within >p90 cortical SBBs (~35%) in comparison to the proportion of these spinules that emerge from boutons/axons in rat CA1 (~12%). Thus, it will be interesting to investigate the origins of spinules within SBBs in CA1 and determine whether activity (e.g. LTP) differentially influences the types of neurites embedded within hippocampal SBBs.

Largely due to evidence demonstrating that increases in electrical and chemical LTP protocols are able to transiently increase the emergence of spinules from dendritic spines (Schuster et al., 1990; Geinisman et al., 1994; Toni et al., 1999; Harris et al., 2003; Ueda and Hayashi, 2013), it has been suggested that spinules function as circuit remodeling and signaling elements and/or as a mechanism to retrieve presynaptic membrane during periods of heightened activity (Spacek and Harris, 2004; Tao-Cheng et al., 2009). While the present study did not directly address spinule function or manipulate activity state, we have demonstrated that cortical boutons progressively engulf spinules over postnatal maturation, but not following the onset of sensory activity or during a period of heightened plasticity in cortex. While it is possible that transient increases in spinule emergence occur immediately following eye-opening and during heightened states of plasticity in cortex, excitatory synaptic boutons do not appear to engulf higher percentages of spinules during these developmental time points. Indeed, if spinules enhance or augment synaptic communication, presynaptic membrane retrieval, or help to maintain synaptic strength/stability, these functions likely depend on spinules first becoming enveloped by presynaptic boutons. Thus, in light of the current developmental and activity-dependent data on spinule emergence and engulfment, we argue for a model of spinule function wherein spinules: (1) act to progressively ‘anchor’ pre- and postsynaptic synaptic elements together over development, potentially ensuring the stability and strength of key synapses within a functional circuit; and (2) enable circuit remodeling and/or membrane retrieval during brief periods of heightened activity.

In addition, it seems likely that the dramatic increase in membrane interface between boutons and spinules within SBBs (Calverley and Jones, 1990; Jones and Calverley, 1991; Rodriguez-Moreno et al., 2018) allows for some form of neuronal communication. For example, following increased activity in hippocampus, a majority of the tips of spinules from postsynaptic spines are coated in clathrin, and these clathrin-coated tips may pinch-off into their enveloping boutons (Spacek and Harris, 2004; Tao-Cheng et al., 2009). Furthermore, the large spinule-bouton membrane interface at synapses might allow for neuronal communication through ephaptic coupling, wherein a spinule that is tightly bound (i.e. high extracellular impedance) to its presynaptic bouton and receives current flow during a presynaptic action potential (Wagner and Djamgoz, 1993; Faber and Pereda, 2018). If these subcellular processes take place, our data on spinule origins suggest that this form of ephaptic coupling would allow for unique forms of communication (e.g. non-synaptic axoaxonic communication) in L4 of V1. Nevertheless, it seems likely that spinules from specific sources (e.g. postsynaptic spines vs. adjacent axons/boutons), and even spinules of different sizes from a single neurite source (Zaccard et al., 2020), will have distinct functional relationships with SBBs. Our data identify spinules from postsynaptic dendritic spines and adjacent axons as prime candidates for investigations into the functional consequences of spinule engulfment by SBBs in the neocortex.

In sum, spinules are conserved structural features of synapses that are embedded within nearly one-quarter of late adolescent excitatory boutons in L4 of V1. Spinules are progressively engulfed by presynaptic boutons as physiological and morphological measures of plasticity wane, suggesting that they have a role in the stabilizing and strengthening a select set of cortical synapses. Determining whether specific subsets of cortical excitatory and inhibitory (Tao-Cheng et al., 2009) boutons preferentially engulf spinules, as well as further defining the mechanisms for spinule induction, expression (Zaccard et al., 2020), maintenance (Ueda and Hayashi, 2013), engulfment, and potential communication, will be crucial to elucidating the role that synaptic spinules play within neuronal microcircuits.

## Acknowledgements

We thank Dr. Claudia Lopez and Dr. Jessica Riesterer from the Multiscale Microscopy Core at OHSU for their technical assistance, with technical support from the OHSU Center for Spatial Systems Biomedicine.

## References Cited

Ahmed B, Anderson JC, Douglas RJ, Martin KA, Nelson JC (1994) Polyneuronal innervation of spiny stellate neurons in cat visual cortex. J Comp Neurol 341:39–49.

Araya R, Vogels TP, Yuste R (2014) Activity-dependent dendritic spine neck changes are correlated with synaptic strength. Proc Natl Acad Sci U S A 111:E2895–2904.

Bailey CH, Thompson EB, Castellucci VF, Kandel ER (1979) Ultrastructure of the synapses of sensory neurons that mediate the gill-withdrawal reflex in Aplysia. Journal of Neurocytology 8:415–444.

Bopp R, Holler-Rickauer S, Martin KA, Schuhknecht GF (2017) An Ultrastructural Study of the Thalamic Input to Layer 4 of Primary Motor and Primary Somatosensory Cortex in the Mouse. J Neurosci 37:2435–2448.

Bozhilova-Pastirova A, Ovtscharoff W (1999) Intramembranous strucutre of synaptic membranes with special reference to spinules in the rat sensorimotor cortex. Eur J Neurosci 11:1843–1846.

Branco T, Marra V, Staras K (2010) Examining size-strength relationships at hippocampal synapses using an ultrastructural measurement of synaptic release probability. Journal of structural biology 172:203–210.

Calverley RKS, Jones DG (1990) Contributions of dendritic spines and perforated synapses to synaptic plasticity. Brain Res Rev 15:215–249.

Case NM, Gray EG, Young JZ (1972) Ultrastructure and synaptic relations in the optic lobe of the brain of Eledone and Octopus. Journa of Ultrastructure Research 39:115–123.

Cheetham CE, Fox K (2010) Presynaptic development at L4 to l2/3 excitatory synapses follows different time courses in visual and somatosensory cortex. J Neurosci 30:12566–12571.

Coleman JE, Nahmani M, Gavornik JP, Haslinger R, Heynen AJ, Erisir A, Bear MF (2010) Rapid structural remodeling of thalamocortical synapses parallels experience-dependent functional plasticity in mouse primary visual cortex. J Neurosci 30:9670–9682.

Colonnier M, Beaulieu C (1985) An empirical assessment of stereological formulae applied to the counting of synaptic disks in the cerebral cortex. J Comp Neurol 231:175–179.

da Costa NM, Martin KA (2009) The proportion of synapses formed by the axons of the lateral geniculate nucleus in layer 4 of area 17 of the cat. J Comp Neurol 516:264–276.

DeFelipe J, Marco P, Busturia I, Merchan-Perez A (1999) Estimation of the number of synapses in the cerebral cortex: Methdological Considerations. Cereb Cortex 9:722–732.

Dufour A, Rollenhagen A, Satzler K, Lubke JHR (2016) Development of Synaptic Boutons in Layer 4 of the Barrel Field of the Rat Somatosensory Cortex: A Quantitative Analysis. Cereb Cortex 26:838–854.

Durack JC, Katz LC (1996) Development of horizontal projections in layer 2/3 of ferret visual cortex. Cereb Cortex 6:178–183.

Erisir A, Dreusicke M (2005) Quantitative morphology and postsynaptic targets of thalamocortical axons in critical period and adult ferret visual cortex. J Comp Neurol 485:11–31.

Faber DS, Pereda AE (2018) Two Forms of Electrical Transmission Between Neurons. Front Mol Neurosci 11:427.

Fiala JC (2005) Reconstruct: a free editor for serial section microscopy. Journal of Microscopy 218:52–61.

Ganeshina O, Berry RW, Petralia RS, Nicholson DA, Geinisman Y (2004) Synapses with a segmented, completely partitioned postsynaptic density express more AMPA receptors than other axospinous synaptic junctions. Neuroscience 125:615–623.

Garcia-Marin V, Kelly JG, Hawken MJ (2019) Major Feedforward Thalamic Input Into Layer 4C of Primary Visual Cortex in Primate. Cereb Cortex 29:134–149.

Gardner MJ, Altman DG (1986) Confidence intervals rather than P values: estimation rather than hypothesis testing. British Medical Journal 292:746–750.

Geinisman Y, Morrell F, deToledo-Morrell L (1987) Axospinous synapses with segmented postsynaptic densities: a morphologically distinct synaptic subtype contributing to the number of profiles of ‘perforated’ synapes visualized in random sections. Brain Res 423:179–188.

Geinisman Y, deToledo-Morrell L, Morrell F (1994) Comparison of Structural Synaptic Modifications Induced by Long-Term Potentiation in the Hippocampal Dentate Gyrus of Young Adult and Aged Rats. Annals of the New York Academy of Sciences 747:452–466.

Geinisman Y, Detoledo-Morrell L, Morrell F, Heller RE, Rossi M, Parshall RF (1993) Structural synaptic correlate of long-term potentiation: Formation of axospinous synapses with multiple, completely partitioned transmission zones. Hippocampus 3:435–445.

Gray EG (1959) Axo-somatic and axo-dendrritic synapses of the cerebral cortex: An electron microscope study. J Anat 93:420–433.

Harris KM, Stevens JK (1989) Dendritic Spines of CA1 Pyramidal Cells in the Rat Hippocampus: Serial Electron Microscopy with Reference to Their Biophysical Characteristics. Journal of Neuroscience 9:2982–2997.

Harris KM, Fiala JC, Ostroff L (2003) Structural changes at dendritic spine synapses during long-term potentiation. Philos Trans R Soc Lond B Biol Sci 358:745–748.

Holderith N, Lorincz A, Katona G, Rozsa B, Kulik A, Watanabe M, Nusser Z (2012) Release probability of hippocampal glutamatergic terminals scales with the size of the active zone. Nat Neurosci 15:988–997.

Issa NP, Trachtenberg JT, Chapman B, Zahs KR, Stryker MP (1999) The critical period for ocular dominance plasticity in the Ferret’s visual cortex. J Neurosci 19:6965–6978.

Jones DG, Calverley RK (1991) Perforated and non-perforated synapses in rat neocortex: three-dimensional reconstructions. Brain Res 556:247–258.

Jones DG, Itarat W, Calverley RK (1991) Perforated synapses and plasticity. A developmental overview. Mol Neurobiol 5:217–228.

Korogod N, Petersen CC, Knott GW (2015) Ultrastructural analysis of adult mouse neocortex comparing aldehyde perfusion with cryo fixation. Elife 4.

Latawiec D, Martin KA, Meskenaite V (2000) Termination of the geniculocortical projection in the striate cortex of macaque monkey: a quantitative immunoelectron microscopic study. J Comp Neurol 419:306–319.

Law MI, Zahs KR, Stryker MP (1988) Organization of primary visual cortex (area 17) in the ferret. J Comp Neurol 278:157–180.

Li Y, Fitzpatrick D, White LE (2006) The development of direction selectivity in ferret visual cortex requires early visual experience. Nat Neurosci 9:676–681.

Liewald JF, Brauner M, Stephens GJ, Bouhours M, Schultheis C, Zhen M, Gottschalk A (2008) Optogenetic analysis of synaptic function. Nat Methods 5:895–902.

Liu X, Ramirez S, Pang PT, Puryear CB, Govindarajan A, Deisseroth K, Tonegawa S (2012) Optogenetic stimulation of a hippocampal engram activates fear memory recall. Nature 484:381–385.

Maffei A, Lambo ME, Turrigiano GG (2010) Critical period for inhibitory plasticity in rodent binocular V1. J Neurosci 30:3304–3309.

Marty S, Peschanski M (1994) Fine structural alteration in target-deprived axonal terminals in the rat thalamus. Neuroscience 62:1121–1132.

Meyer D, Bonhoeffer T, Scheuss V (2014) Balance and stability of synaptic structures during synaptic plasticity. Neuron 82:430–443.

Murphy DD, Andrews SB (2000) Culture models for the study of estradiol-induced synaptic plasticity. Journal of Neurocytology 29:411–417.

Murthy VN, Schikorski T, Stevens CF, Zhu Y (2001) Inactivity produces increases in neurotransmitter release and synapse size. Neuron 32:673–682.

Ostroff LE, Fiala JC, Allwardt B, Harris KM (2002) Polyribosomes redistribute from dendritic shafts into spines with enlarged synapses during LTP in developing rat hippocampal slices. Neuron 35:535–545.

Pappas G, Purpura D (1961) Fine Structure of dendrites in the superficial neocortical neuropil. Exp Neurol 4:507–530.

Peters A, Payne BR, Budd J (1994) A numerical analysis of the geniculocortical input to striate cortex in the monkey. Cereb Cortex 4:215–229.

Petralia RS, Wang YX, Mattson MP, Yao PJ (2015) Structure, Distribution, and Function of Neuronal/Synaptic Spinules and Related Invaginating Projections. Neuromolecular medicine 17:211–240.

Petralia RS, Wang YX, Mattson MP, Yao PJ (2018) Invaginating Structures in Mammalian Synapses. Frontiers in synaptic neuroscience 10:4.

Qiao Q, Ma L, Li W, Tsai JW, Yang G, Gan WB (2016) Long-term stability of axonal boutons in the mouse barrel cortex. Dev Neurobiol 76:252–261.

Quinn DP, Kolar A, Harris SA, Wigerius M, Fawcett JP, Krueger SR (2019) The Stability of Glutamatergic Synapses Is Independent of Activity Level, but Predicted by Synapse Size. Frontiers in cellular neuroscience 13:291.

Rodriguez-Moreno J, Rollenhagen A, Arlandis J, Santuy A, Merchan-Perez A, DeFelipe J, Lubke JHR, Clasca F (2018) Quantitative 3D Ultrastructure of Thalamocortical Synapses from the “Lemniscal” Ventral Posteromedial Nucleus in Mouse Barrel Cortex. Cereb Cortex 28:3159–3175.

Schindelin J, Arganda-Carreras I, Frise E, Kaynig V, Longair M, Pietzsch T, Preibisch S, Rueden C, Saalfeld S, Schmid B, Tinevez J-Y, White DJ, Hartenstein V, Eliceiri K, Tomancak P, Cardona A (2012) Fiji: an open-source platform for biological-image analysis. Nature Methods 9:676–682.

Schuster T, Krug M, Wenzel J (1990) Spinules in axospinous synapses of the rat dentate gyrus: changes in density following long-term potentiation. Brain Research 523:171–174.

Sherrington C (1906) The integrative action of the nervous system. New Haven, CT: Yale University Press.

Smith GB, Sederberg A, Elyada YM, Van Hooser SD, Kaschube M, Fitzpatrick D (2015) The development of cortical circuits for motion discrimination. Nat Neurosci 18:252–261.

Sorra KE, Fiala JC, Harris KM (1998) Critical assessment of the involvement of perforations, spinules, and spine branching in hippocampal synpase formation. J Comp Neurol 398:225–240.

Spacek J (1985) Relationships between synaptic junctions, puncta adherentia and the spine apparatus at neocortical axo-spinous synapses. A serial section study. Anat Embryol (Berl) 173:129–135.

Spacek J, Harris KM (2004) Trans-endocytosis via spinules in adult rat hippocampus. J Neurosci 24:4233–4241.

Tao-Cheng JH, Dosemeci A, Gallant PE, Miller S, Galbraith JA, Winters CA, Azzam R, Reese TS (2009) Rapid turnover of spinules at synaptic terminals. Neuroscience 160:42–50.

Tarrant SB, Routtenberg A (1977) The synaptic spinule in the dendritic spine: electron microscopic study of the hippocampal dentate gyrus. Tissue Cell 9:461–473.

Toni N, Buchs PA, Nikonenko I, Bron CR, Muller D (1999) LTP promotes formation of multiple spine synapses between a single axon terminal and a dendrite. Nature 402:421–425.

Ueda Y, Hayashi Y (2013) PIP(3) regulates spinule formation in dendritic spines during structural long-term potentiation. J Neurosci 33:11040–11047.

Wagner H, Djamgoz M (1993) Spinules: a case for retinal synaptic plasticity. Trends Neurosci 16:201–206.

Westrum LE, Blackstad TW (1962) An electron microscopic study of the stratum radiatum of the rat hippocampus (regio superior, CA 1) with particular emphasis on synaptology. J Comp Neurol 119:281–309.

Wu Y, Whiteus C, Xu CS, Hayworth KJ, Weinberg RJ, Hess HF, De Camilli P (2017) Contacts between the endoplasmic reticulum and other membranes in neurons. Proc Natl Acad Sci U S A 114:E4859–E4867.

Yu H, Majewska AK, Sur M (2011) Rapid experience-dependent plasticity of synapse function and structure in ferret visual cortex in vivo. Proc Natl Acad Sci U S A 108:21235–21240.

Zaccard CR, Shapiro L, Martin-de-Saavedra MD, Pratt C, Myczek K, Song A, Forrest MP, Penzes P (2020) Rapid 3D Enhanced Resolution Microscopy Reveals Diversity in Dendritic Spinule Dynamics, Regulation, and Function. Neuron.

